# Bioengineering secreted proteases converts divergent Rcr3 orthologs and paralogs into extracellular immune co-receptors

**DOI:** 10.1101/2024.02.14.580413

**Authors:** Jiorgos Kourelis, Mariana Schuster, Fatih Demir, Oliver Mattinson, Sonja Krauter, Parvinderdeep S. Kahlon, Ruby O’Grady, Samantha Royston, Ana Lucía Bravo-Cazar, Brian C. Mooney, Pitter F. Huesgen, Sophien Kamoun, Renier A. L. van der Hoorn

**Affiliations:** The Plant Chemetics Laboratory, Department of Biology, University of Oxford, South Parks Road, OX1 3RB Oxford, UK; The Sainsbury Laboratory, Norwich Research Park, NR4 7UH, Norwich, UK; Central Institute for Engineering, Electronics and Analytics (ZEA), Analytics (ZEA-3), Research Centre Jülich, Wilhelm-Johnen-Str., 52428 Jülich, Germany; Laboratory of Plant Physiology, Plant Sciences Group, Wageningen University & Research, Droevendaalsesteeg 1, 6708PB, Wageningen, The Netherlands; Department of Life Sciences, Imperial College, SW7 2AZ, London, UK; Leibniz Institute of Plant Biochemistry, Weinberg 3, 06120, Halle, Germany; Department of Biomedicine, Aarhus University, Høegh-Guldbergsgade 10, DK-8000 Aarhus; Faculty of Biology, University of Freiburg, Schänzlestrasse 1, 79104 Freiburg, Germany

**Keywords:** tomato, plant immunity, *Cladosporium fulvum*, hypersensitive response, disease resistance, protease engineering

## Abstract

Secreted immune proteases Rcr3 and Pip1 of tomato are both inhibited by Avr2 from the fungal plant pathogen *Cladosporium fulvum* but only Rcr3 act as a decoy co-receptor that detects Avr2 in the presence of the Cf-2 immune receptor. Here, we identified crucial residues from tomato Rcr3 required for Cf-2-mediated signalling and bioengineered various proteases to trigger Avr2/Cf-2 dependent immunity. Despite substantial divergences in Rcr3 orthologs from eggplant and tobacco, only minimal alterations were sufficient to trigger Avr2/Cf-2-triggered immune signalling. Tomato Pip1, by contrast, was bioengineered with 16 Rcr3-specific residues to initiate Avr2/Cf-2-triggered immune signalling. These residues cluster on one side next to the substrate binding groove, indicating a potential Cf-2 interaction site. Our findings also revealed that Rcr3 and Pip1 have distinct substrate preferences determined by two variant residues and that both are suboptimal for binding Avr2. This study advances our understanding of Avr2 perception and opens avenues to bioengineer proteases to broaden pathogen recognition in other crops.

## INTRODUCTION

The fungal pathogen *Cladosporium fulvum* (syn. *Passalora fulva*) causes leaf mould disease in tomato (*Solanum lycopersicum* L.). While most cloned resistance (*R)* genes of plants are nucleotide-binding oligomerization domain (NOD)-like receptors (NLRs), all cloned *R* genes against *C. fulvum* encode for cell-surface localized receptor-like proteins (RLPs) with extracellular leucine-rich repeats (LRRs). One such *R* gene encoding a LRR-RLP is *Cf-2*, which was introgressed into cultivated tomato (*Solanum lycopersicum*) from currant tomato (*S. pimpinellifolium*) (Dixon et al., 1996). *Cf-2* confers recognition of the secreted small cysteine-rich effector Avr2 of *C. fulvum* (Luderer et al., 2002), ultimately resulting in a localized programmed cell death called the hypersensitive response (HR) and stopping further pathogen growth.

Recognition of Avr2 by Cf-2 requires tomato *Rcr3*, which encodes a papain-like cysteine protease (PLCP; Krüger et al., 2002). PLCPs are stable, 20-25 kDa endopeptidases with a catalytic Cys-His-Asn triad that contains the catalytic Cys residue in a substrate binding groove (Shindo and van der Hoorn, 2008). PLCPs are produced as pre-pro-proteases that have a signal peptide for secretion and remain inactive until the autoinhibitory prodomain is removed. The mature protease domain of Rcr3 is sufficient to trigger Avr2/Cf-2-dependent HR (Rooney et al., 2005) and Rcr3 activation in tomato is promoted by the highly abundant apoplastic subtilase P69B (Paulus et al., 2020).

Avr2 binds and inhibits mature Rcr3 and is not proteolytically processed by Rcr3 (Rooney et al., 2005). *Cf-2* tomato lines lacking *Rcr3* do not display obvious phenotypes (Dixon et al., 2000), and catalytically inactive Rcr3 can still bind Avr2 and trigger Cf-2-dependent HR (Paulus et al., 2020). These two facts indicate that it is not the substrate/product of Rcr3 that is recognized by Cf-2, but rather the Avr2/Rcr3 complex itself. Further studies using mutant Avr2 protein showed that the inhibition of Rcr3 by Avr2 positively correlates with the induction of Cf-2-dependent HR (van’t Klooster et al., 2011), and that Rcr3 homologs with the naturally occurring N194D substitution cannot be inhibited by Avr2, and do not trigger Avr2/Cf-2-dependent HR (Shabab et al., 2008; Hörger et al., 2012). Although these data all indicate that Cf-2 physically interacts with the Avr2/Rcr3 complex, this interaction remains to be demonstrated.

The indirect perception of Avr2 is consistent with the guard hypothesis, which proposes that immune receptors ‘guard’ the virulence target of pathogen-derived effector proteins, rather than the effector itself (van der Biezen and Jones, 1998; van der Hoorn et al., 2002; Dangl and Jones, 2001). Accordingly, Cf-2 ‘guards’ Rcr3 to monitor its manipulation by Avr2. Avr2 also inhibits Pip1 (*Phytophthora*-inhibited protease-1; Tian et al., 2007), a paralog of Rcr3 that is encoded by the same genomic locus but is evolving independently from Rcr3 in solanaceous plants (Ilyas et al., 2015; Kourelis et al., 2020). Pip1 is much more abundant than Rcr3, and Pip1 depletion by RNA interference increases susceptibility to *C. fulvum*, revealing that Pip1 is an immune protease and the operative virulence target for Avr2 (Ilyas et al., 2015). By contrast, in the absence of Cf-2, *rcr3* mutant lines are not more susceptible to *C. fulvum* infection, indicating that Rcr3 is not an operative target but acts as a decoy to trap the fungal pathogen into a recognition event in plants carrying Cf-2 (Shabab et al., 2008; van der Hoorn and Kamoun, 2008).

Rcr3 and Pip1 evolved >50 million years ago (Mya) because homologs are present in solanaceous plants that diverged in this period (Kourelis et al., 2020). By contrast, Cf-2 only occurs in the *Solanum* genus which evolved ∼8Mya, indicating that Cf-2 co-opted an existing protease to detect its manipulation during infection (Kourelis et al., 2020). Most *Solanum* Rcr3 homologs can trigger Avr2/Cf-2-dependent HR, but many Rcr3 homologs outside the *Solanum* genus cannot, even though many can be inhibited by Avr2 (Kourelis et al., 2020). This indicates that the residues required for the interaction with Avr2, and those required for *Cf-2*-dependent Avr2 recognition are distinct, and is consistent with the adaptation of Cf-2 to guard Rcr3 in *Solanum* species.

Here we investigated which residues in Rcr3 are required to trigger Avr2/Cf-2-dependent HR. This knowledge is required for future bioengineering of this perception system to mediate recognition of protease inhibitors produced by other plant pathogens and in other (crop) plants. To do this, we bioengineered three Rcr3-like proteases with increasing phylogenetic distance to *Solanum* Rcr3, starting with eggplant Rcr3, then tobacco Rcr3, and finally tomato Pip1. In all these proteases, we were able to bioengineer Avr2/Cf-2-dependent HR, providing intriguing insights into the evolution of this unique perception mechanism and an important platform for future decoy bioengineering.

## RESULTS

### A single residue substitution in eggplant SmRcr3 is sufficient for Avr2/Cf-2-dependent HR

While tomato and potato Rcr3 homologs can complement Cf-2/Avr2-dependent HR in *N. benthamiana*, the eggplant (*Solanum melongena* L.) Rcr3 (*Sm*Rcr3) cannot trigger Cf-2/Avr2-dependent HR upon agroinfiltration (Kourelis et al., 2020). The mature protease domain of eggplant *Sm*Rcr3 differs at 45 residues from Rcr3 of *S. pimpinellifolium* (*Sp*Rcr3, here referred to as Rcr3, **Figure 1A**). To identify a region in *Sm*Rcr3 that prevents signalling through Cf-2, we replaced three arbitrary, similar-sized parts of *Sm*Rcr3 with the corresponding Rcr3 sequence (**Figure 1B**). *Sm*Rcr3 carrying part-4 of Rcr3 was able to confer Cf-2/Avr2-dependent HR, whilst *Sm*Rcr3 carrying part-5 of Rcr3 causes a very weak HR (**Figure 1C**). part-4 of *Sm*Rcr3 contains 12 variant residues, including one aspartic acid residue, (D244), which is a proline in all Rcr3 homologs that can trigger Cf-2/Avr2-dependent HR (Kourelis et al., 2020). Most other variant residues in part-4 are also different between *Solanum* Rcr3 homologs that can trigger Cf-2/Avr2-dependent HR.

**Figure 1.**
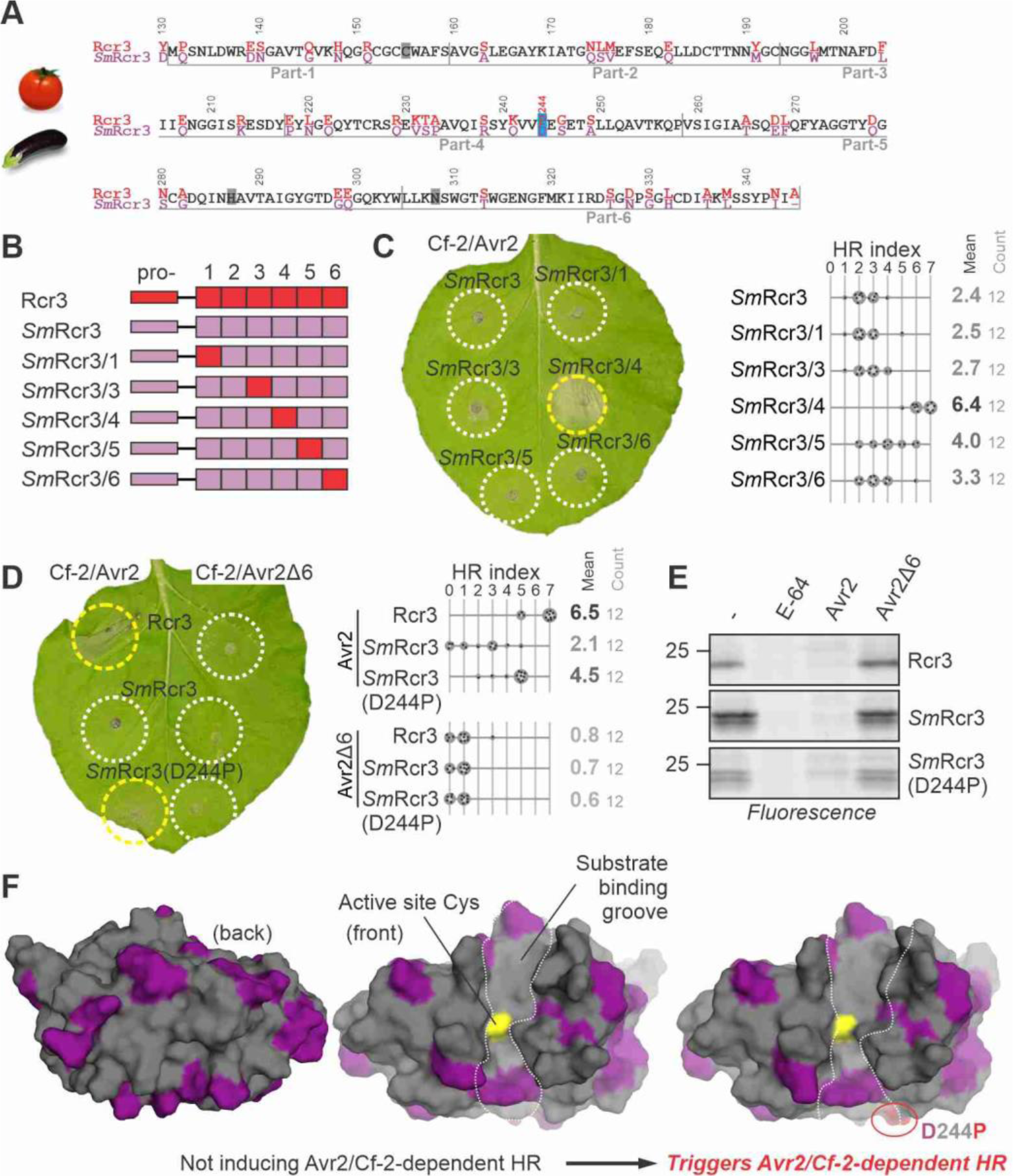
A single residue substitution in eggplant Rcr3 triggers Avr2/Cf-2-dependent HR. **(A)** Alignment of tomato Rcr3 and *Sm*Rcr3 protein sequences. The numbering is based on the Rcr3 sequence of cultivated tomato (*Sl*Rcr3), which carries one additional amino acid in the prodomain compared to Rcr3. **(B)** Tested Rcr3/*Sm*Rcr3 hybrids. The mature protease domain of *Sm*Rcr3 was split into six parts and each part was replaced by the corresponding part of Rcr3. **(C)** *Sm*Rcr3 carrying part-4 of Rcr3 triggers Avr2/Cf2-2-dependent HR. *Sm*Rcr3 and five of its hybrids were transiently co-expressed with Avr2 and Cf-2 in *N. benthamiana*, and the hypersensitive response (HR) was scored 5 days later. Shown is a representative leaf with infiltrated regions encircled with a white line (no HR) or yellow line (HR). The HR index was determined using n=12 replicates. **(D)** *Sm*Rcr3 does not trigger Avr2/Cf2-2-dependent HR, but *Sm*Rcr3(D244P) does. Rcr3, *Sm*Rcr3 and *Sm*Rcr3(D244P) were transiently co-expressed with Avr2/Avr2Δ6 and Cf-2 in *N. benthamiana*, and the hypersensitive response (HR) was scored 5 days later. Shown is a representative leaf with infiltrated regions encircled with a white line (no HR) or yellow line (HR). The HR index was determined using n=12 replicates. **(E)** Rcr3, *Sm*Rcr3 and *Sm*Rcr3(D244P) are active proteases that can be inhibited by Avr2. Apoplastic fluids isolated from agroinfiltrated leaves transiently expressing Rcr3, *Sm*Rcr3 and *Sm*Rcr3(D244P) were pre-incubated for 45 min with and without 100 µM E-64, or 1 µM Avr2 or inactive Avr2Δ6, and then labelled for 3 hours with 0.2 µM MV201. Samples were separated on SDS-PAGE gels and scanned for fluorescence. **(F)** Structural models of *Sm*Rcr3 and bioengineered *Sm*Rcr3. Both sequences were modelled using AlphaFold2 (both pTM= 0.94), and presented in PyMol using surface representation of the front of the protease with residues that are identical to tomato Rcr3 (grey) and that are specific to *Sm*Rcr3 (purple). Bioengineering of *Sm*Rcr3 with D224P (red) causes Avr2/Cf-2-dependent HR.

*Sm*Rcr3 with the D244P mutation indeed triggers HR upon co-expression with Cf-2 and Avr2 (**Figure 1D**). Furthermore, *Sm*Rcr3 produced upon agroinfiltration can be labelled with MV201 (**Figure 1E**; Kourelis et al., 2020), a fluorescent activity-based probe for PLCPs (Richau et al., 2012), and preincubation with Avr2 prevents *Sm*Rcr3 labelling by MV201 (**Figure 1E**). This confirm that *Sm*Rcr3 is an active protease that can be inhibited by Avr2. This inhibition is specific, as preincubation with Avr2 lacking the last six residues (Avr2Δ6), an inactive protease inhibitor (van’t Klooster et al., 2011), does not prevent *Sm*Rcr3 labelling by MV201 (**Figure 1E**). This experiment shows that a single D244P substitution is sufficient to bioengineer eggplant *Sm*Rcr3 into a protein that triggers Avr2/Cf-2-dependent HR, and that interaction with Avr2 and with Cf-2 can be uncoupled in Rcr3 homologs.

We summarised the predicted location of the identified residues required for Avr2/Cf-2-dependent HR by creating structural models for *Sm*Rcr3 and the bioengineered *Sm*Rcr3(D224P) proteases using AlfaFold2 (Jumper et al., 2021). The structure of Avr2 could not be predicted by Alphafold2, probably because of a shallow multi sequence alignment (MSA). By contrast, there is a robust MSA for the proteases. There are many well-resolved crystal structures for papain-like cysteine proteases and the structures are predicted with a high predicted Template Modeling score (pTM=0.94). As with all PLCPs, all protease models consist of two lobes with the substrate binding groove in between. The catalytic Cys is located in the middle of that substrate binding groove at the end of a long alpha helix, and is in close proximity to the catalytic His and Asn residues. Residues in *Sm*Rcr3 that differ from Rcr3 scatter all over the front and back surfaces of these proteases (**Figure 1F**, purple) although the substrate binding groove is relatively similar. Most of these variant residues, however, are not required for Avr2/Cf-2-dependent HR. The D244P substitution locates at the edge of the substrate binding groove and is predicted to have minimal effect on the local structure of the protease (**Figure 1F**), although the effect of missense mutations may be difficult to predict by AlphaFold2 (Buel & Walters, 2022). Although *Sm*Rcr3(D244P) is not as active in triggering Avr2/Cf-2-dependent HR as tomato Rcr3, the fact that HR can be triggered with a single substitution in a protease that is so different from tomato Rcr3 is remarkable.

### G194N substitution and a 3-residue insertion in Nicotiana Rcr3 triggers Avr2/Cf-2-dependent HR

We previously found that *Nicotiana* Rcr3 homologs also cannot trigger Avr2/Cf-2-dependent HR (Kourelis et al., 2020). *Nicotiana* Rcr3 homologs differ at 49 residues and a 3-amino acid deletion in the protease domain compared to Rcr3 (**Figure 2A**). While *N. benthamiana* lacks a functional Rcr3 homolog, we previously found that the inactivated *NbRcr3a* gene can be ‘resurrected’ as an active protease, called r*Nb*Rcr3 (Kourelis et al., 2020). However, this r*Nb*Rcr3 cannot be inhibited by Avr2, probably because it lacks a key residue required for Avr2 inhibition (N194; Shabab et al., 2008; Hörger et al., 2012), which is a glycine in r*Nb*Rcr3.

**Figure 2.**
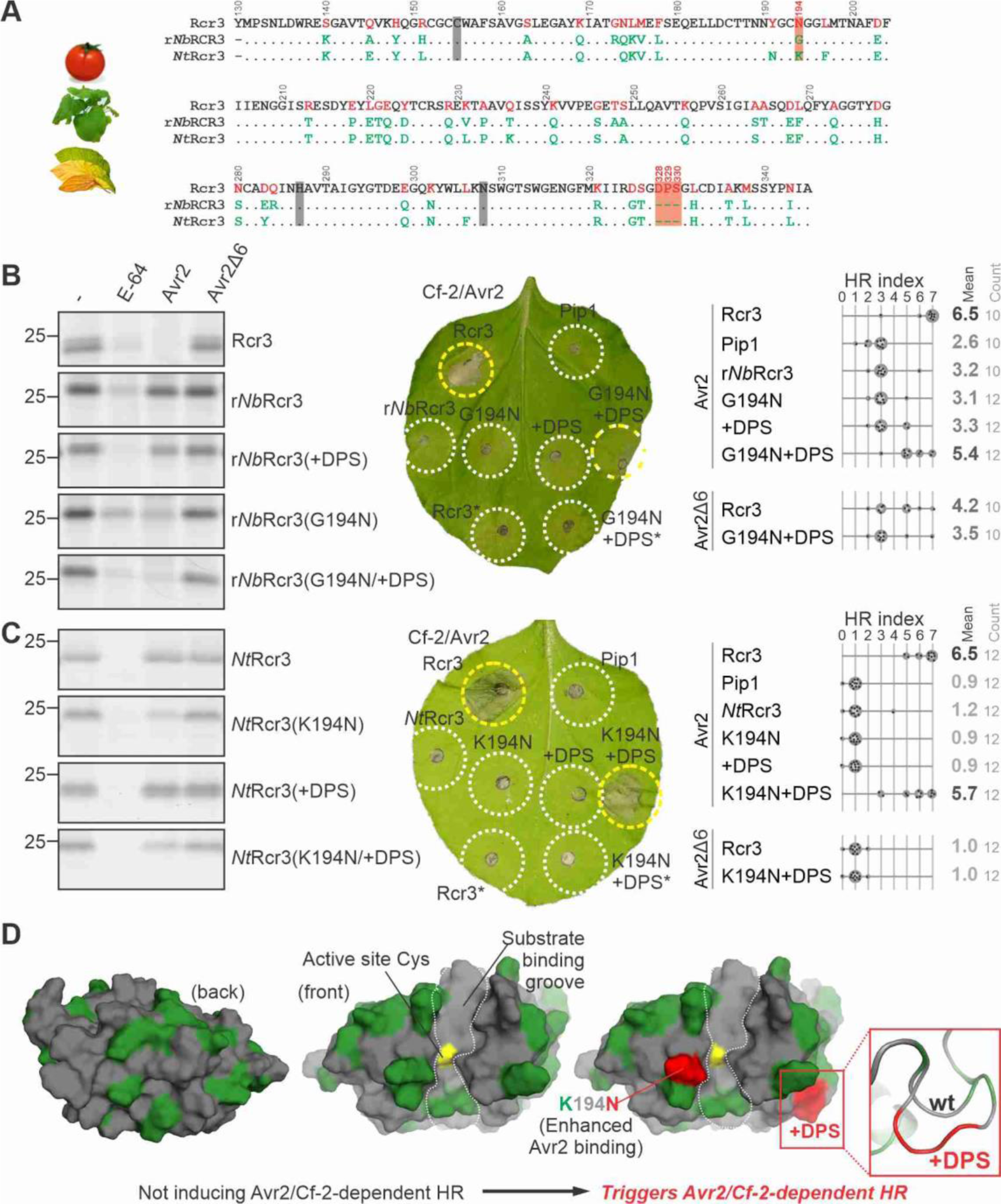
N194 and a 3-residue insertion in *Nicotiana* Rcr3 triggers Avr2/Cf-2-dependent HR. **(A)** Alignment of tomato Rcr3 with Rcr3 homologs from *N. benthamiana* (*Nb*) and *N. tabacum* (*Nt*). The numbering is based on the *Sl*Rcr3 sequence. **(B)** G194N substitution in resurrected *Nb*Rcr3 (r*Nb*Rcr3) reconstitutes Avr2 inhibition (left) and an additional 3-residue insertion (+DPS) triggers Avr2/Cf-2-dependent HR (left). **(C)** K194N substitution in tobacco *Nt*Rcr3 reconstitutes Avr2 inhibition (left) and an additional 3-residue insertion (+DPS) triggers Avr2/Cf-2-dependent HR (left). **(B-C)** Left gel images: Apoplastic fluids isolated from agroinfiltrated leaves transiently expressing (mutant) Rcr3 (homologs) were pre-incubated for 45 min with and without 100 µM E-64, or 1 µM Avr2 or inactive Avr2Δ6, and then labelled for 3 hours with 0.2 µM fluorescent activity-based probe MV201 to detect the proteases that are not inhibited. Samples were separated on SDS-PAGE gels and scanned for fluorescence. Right HR assays: (mutant) Rcr3 homologs and Pip1 were transiently co-expressed with Avr2/Avr2Δ6 and Cf-2 in *N. benthamiana*, and the hypersensitive response (HR) was scored 5 days later. Shown is a representative leaf with infiltrated regions encircled with a white line (no HR), yellow line (HR), or both (intermediate HR). *, co-expressed with /Avr2Δ6 instead of Avr2. The HR index was determined using n=12 replicates. **(D)** Structural models of *Nt*Rcr3 and bioengineered *Nt*Rcr3. Both sequences were modelled using AlphaFold2 (both pTM= 0.94), and presented in PyMol using surface representation of the front of the protease with residues that are identical to tomato Rcr3 (grey) and that are specific to *Nt*Rcr3 (green). Bioengineering of *Nt*Rcr3 with K194N and inserting tripeptide DPS (red) causes Avr2/Cf-2-dependent HR.

To test if the G194N substitution in r*Nb*Rcr3 promotes sensitivity for Avr2 inhibition, we preincubated r*Nb*Rcr3 and r*Nb*Rcr3(G194N) with Avr2, E-64, or Avr2Δ6 (van’t Klooster et al., 2011) and labelled the remaining active protease with activity-based probe MV201. Detection by scanning the proteins upon separating on proteins gels for fluorescence showed that r*Nb*Rcr3 carrying the G194N substitution can be inhibited by Avr2 (**Figure 2B**). However, despite being Avr2 sensitive, co-expression with Avr2 and Cf-2 showed that r*Nb*Rcr3(G194N) does still not trigger Avr2/Cf-2-dependent HR (**Figure 2B** and Supplemental **Figure S1**).

All *Nicotiana* Rcr3 homologs lack three residues (D328/P329/S330) compared to Rcr3 homologs of *Solanum* and *Capsicum* (Kourelis et al., 2020). r*Nb*Rcr3 carrying G194N and these three additional residues (r*Nb*Rcr3(G194N+DPS)) is an active protease, sensitive for Avr2 inhibition and also triggers Avr2/Cf-2-dependent HR (**Figure 2B** and Supplemental **Figure S1**). By contrast, r*Nb*Rcr3(+DPS) is unable to interact with Avr2 and does not trigger Avr2/Cf-2-dependent HR (**Figure 2B** and Supplemental **Figure S1**). These data demonstrate that the DPS insertion is required for HR induction but not for Avr2 inhibition.

Like r*Nb*Rcr3, *N. tabacum* Rcr3 (*Nt*Rcr3) also lacks the DPS insertion and carries an N194 substitution, in this case into a lysine (K). As with r*Nb*Rcr3, Avr2 inhibition can be bioengineered into *Nt*Rcr3 with the K194N substitution, but these proteases do not trigger Avr2/Cf-2-dependent HR (**Figure 2C**). The additional DPS insertion is also required to trigger Avr2/Cf-2-dependent HR (**Figure 2C**). *Nt*Rcr3 carrying only the DPS insertion is not inhibited by Avr2 and is unable to trigger HR (**Figure 2C**). These data demonstrate again that the DPS insertion is required for HR induction in *Nicotiana* Rcr3 homologs but not for Avr2 inhibition.

Modelling the structure of both *Nt*Rcr3 and the engineered *Nt*Rcr3 with AlphaFold2 resulted in reliable complexes (pTM = 0.94), that show that the variant residues in *Nt*Rcr3 when compared to tomato Rcr3 scatter over the surface on both sides of the protein (**Figure 2D**, green residues). The K194N substitution is next to the catalytic Cys residue at the edge of the substrate binding groove. This substitution promotes Avr2 binding, suggesting that Avr2 would interact with this residue and occupies the active site, consistent with its ability to suppress labeling with activity-based probe MV201, which labels the active site Cys residue. The insertion of the DPS tripeptide (D328/P329/S330) into *Nt*Rcr3 is predicted to extend a loop on the side of the protease (**Figure 2D**). This DPS tripeptide locates in a similar region as the D224P mutation in *Sm*Rcr3, although in an adjacent loop. This indicates that the residues required for Avr2/Cf-2-dependent HR are clustered on the right lobe of the Rcr3 protease in both *Sm*Rcr3 and *Nt*Rcr3.

### Pip1 triggers Avr2/Cf-2-dependent HR when carrying three parts of Rcr3

Next, we bioengineered *Solanum lycopersicum* Pip1 (*Sl*Pip1, here referred to as Pip1), a tomato immune protease that is even more distantly related to its paralog Rcr3 than *Sm*Rcr3 and *Nt*Rcr3 (Ilyas et al., 2015). The mature protease domains of Rcr3 and Pip1 differ at 93 residues, with several amino acid insertions and deletions (**Figure 3A**). Pip1 can be inhibited by Avr2 (Shabab et al., 2008), but it does not trigger Avr2/Cf-2-dependent HR (Kourelis et al., 2020). One notable difference is that the DPS amino acid sequence in Rcr3, which is absent from *Nicotiana* Rcr3 and essential for Avr2/Cf-2-dependent HR, corresponds to a VDG sequence in Pip1 (**Figure 3A**). Interestingly, this sequence is VHG in *S. pimpinellifolium* Pip1 (*Sp*Pip1) which has been co-introgressed with Rcr3 when creating the MM-Cf2 line. The D329H variant residue changing VDG into VHG is the only difference between mature Pip1 of cultivated tomato and *Sp*Pip1 of *S. pimpinellifolium*.

**Figure 3.**
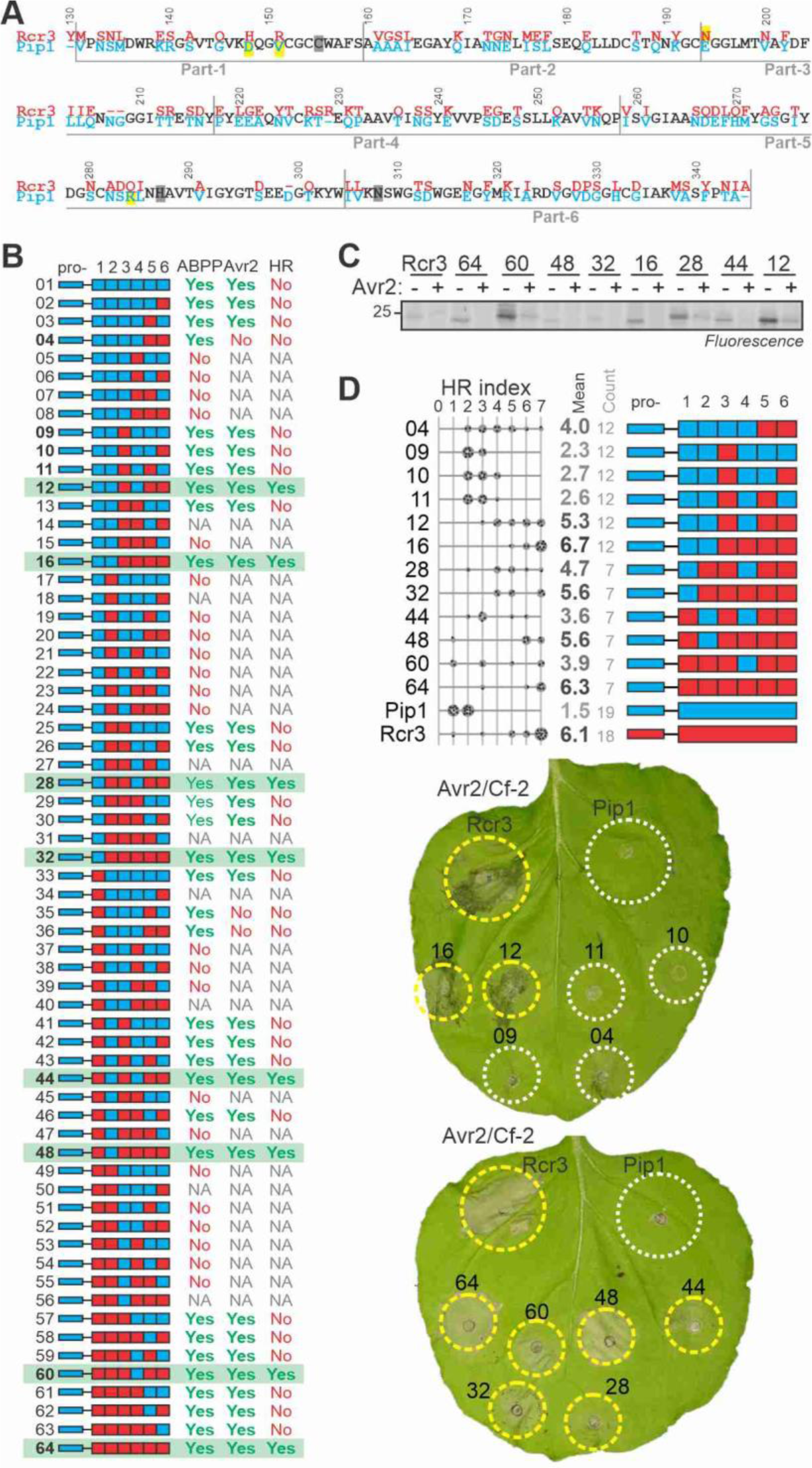
Pip1 triggers Avr2/Cf-2-dependent HR when carrying three parts of Rcr3. **(A)** Amino acid alignment if Pip1 and Rcr3. The numbering is based on the Rcr3 sequence. Four residues contributing to Avr2 inhibition are highlighted in yellow. **(B)** Summary of analysed Pip1/Rcr3 hybrids, showing construct number (pJK); domain architecture; labelling of the protease with MV201; inhibition by Avr2; and Avr2/Cf-2-dependent HR. NA, not analysed. The eight Pip1/Rcr3 hybrids that trigger HR are highlighted in green. **(C)** Eight HR-inducing Pip1/Rcr3 hybrids are active proteases that can be inhibited by Avr2. Apoplastic fluids isolated from agroinfiltrated leaves transiently expressing Pip1/Rcr3 hybrids were pre-incubated for 45 min with and without 500 nM Avr2 and then labelled for 5 hours with 0.2 µM MV201. Samples were separated on SDS-PAGE gels and scanned for fluorescence. **(D)** Quantification of HR for selected Pip1/Rcr3 hybrids. Pip1/Rcr3 hybrids were transiently co-expressed with Avr2 and Cf-2 in *N. benthamiana*, and the hypersensitive response (HR) was scored 5 days later. Shown is a representative leaf with infiltrated regions encircled with a white line (no HR) or yellow line (HR). The HR index was determined using n=12 replicates.

To identify the specific Rcr3 residues required for triggering Avr2/Cf-2-dependent HR, we divided Rcr3 into seven arbitrary similarly-sized fragments and replaced corresponding segments in Pip1 with these Rcr3 fragments. As expected, the prodomain of Pip1 fused to the Rcr3 mature protease still triggers Avr2/Cf-2-dependent HR (**Figure 3B**, construct 64), so all our remaining constructs carry the Pip1 prodomain. We next exchanged the six parts of the Pip1 protease domain with the corresponding parts of Rcr3. Out of the 64 possible combinations, we cloned and transiently expressed 57 Pip1/Rcr3 hybrids in *N. benthamiana* by agroinfiltration.

Of the 57 tested hybrid proteases, 33 were active proteases, as determined by MV201 labelling of apoplastic fluids isolated from agroinfiltrated leaves (**Figure 3B** and Supplemental **Figure S2**). Given that all hybrids have an intact catalytic triad, we speculate that the 24 inactive Rcr3/Pip1 hybrids are unstable due to a disruption of the core structure of the protease caused by a mismatch of structural residues in the hybrids. Most unstable hybrids have either part-2 of Rcr3 combined with part-3 of Pip1, or part-3 of Pip1 combined with part-4 of Rcr3 (**Figure 3B**). Of the 33 active proteases, 30 could be inhibited by Avr2 (**Figure 3B** and Supplemental **Figure S3**). The three Avr2-insensitive hybrid proteases contain combinations of residues H148, R151, E194 and Q284, which we previously showed to reduce Avr2 inhibition (Kourelis et al., 2020): construct 04 carries both E194 and Q284, whereas constructs 35 and 36 carry H148, R151, E194 and Q284. Interestingly, because Pip1 carries D148, V151 and R284 and Rcr3 carries only N194, this means that both Pip1 and Rcr3 are suboptimal interactors of Avr2.

Of the 30 Avr2-sensitive hybrid proteases, only eight can trigger *Cf-2*-dependent HR upon co-infiltration with Avr2 into leaflets of tomato MM-*Cf-2 rcr3-3* lines, which lack Rcr3 (**Figure 3B** and Supplemental **Figure S3**). All these HR-inducing hybrids are active proteases that can be inhibited by Avr2 (**Figure 3C**). All the HR-inducing hybrids contain N194 in part-3 but many carry either D148 and V151 in part-1 or R284 in part-5, indicating that not all residues that strengthen the interaction with Avr2 are required for triggering HR. Pip1 carrying three parts from Rcr3 (parts 3, 5 and 6, see construct 12 in **Figure 3B**), can trigger Avr2/Cf-2-dependent HR, which is consistent with the fact that all HR-inducing hybrids carry the same three Rcr3 parts (**Figure 3B**). Replacing any of these Rcr3 parts for the corresponding Pip1 part results in a loss of HR (**Figure 3B**), even though these proteases are active and inhibited by Avr2 (Supplemental **Figures S3 and S4**). These data indicate that Avr2/Cf-2-dependent HR requires multiple residues residing in three of the six parts of the mature protease.

A more detailed, quantitative analysis of HR by co-expression of Pip1/Rcr3 hybrids with Avr2 and Cf-2 in *N. benthamiana* confirms that parts-5 and -6 of Rcr3 are required for inducing Avr2/Cf-2 independent HR, while part-3 makes a significant quantitative contribution (**Figure 3D**). Hybrid 12, containing parts-3, -5 and 06 of Rcr3, can induce HR, whereas hybrids 10 and 11, lacking parts-5 or -6 from Rcr3, respectively, cannot induce HR (**Figure 3D**). Hybrid 04, lacking part-3 of Rcr3, induces HR but to a quantitatively lesser extend (**Figure 3D**). Notably all these hybrids are active proteases that can be inhibited by Avr2 (Supplemental **Figure S2**). Part-4 also contributes to the strength of HR, since constructs containing part-4 of Pip1 have a consistently reduced strength in inducing HR (**Figure 3D**). However, this contribution to HR by part-4 of Rcr3 is relatively weak compared to parts-3, -5, and -6 and part-4 was not investigated further.

### Further mutagenesis identifies Rcr3 residues required for Avr2/Cf-2-dependent HR

To further identify critical residues in Rcr3 required to trigger Cf-2/Avr2-dependent HR, we took the Pip1 hybrid protease containing parts 3, 5, and 6 from Rcr3, and made a series of substitutions in each Rcr3 part. These mutants were analysed by Activity-based Protein Profiling (ABPP) to confirm that they accumulate as active proteases, and co-expression with Avr2 and Cf-2 to determine the relative strength of HR induction.

Part-3 contains 10 variant residues and two deletions and includes the N194 residue which promotes Avr2 binding in Rcr3 when compared to E194 in Pip1 (Kourelis et al., 2020). We tested eleven mutants with various combinations of variant residues. Reflecting the quantitative contribution of part-3, all mutant proteases were able to trigger Avr2/Cf-2-dependent HR, albeit with varying strengths (**Figure 4A**; **Table S1**). Mutants carrying E194 from Pip1 were consistently less able to trigger HR when compared to those carrying N194 of Rcr3 (**Figure 4A**), consistent with N194 promoting the interactions with Avr2 (Shabab et al., 2008; Hörger et al., 2012; Kourelis et al., 2020). No contributions to HR were identified for the other variant residues in part-3. These data indicate that N194 is the main contributor to HR in Rcr3 part-3.

**Figure 4.**
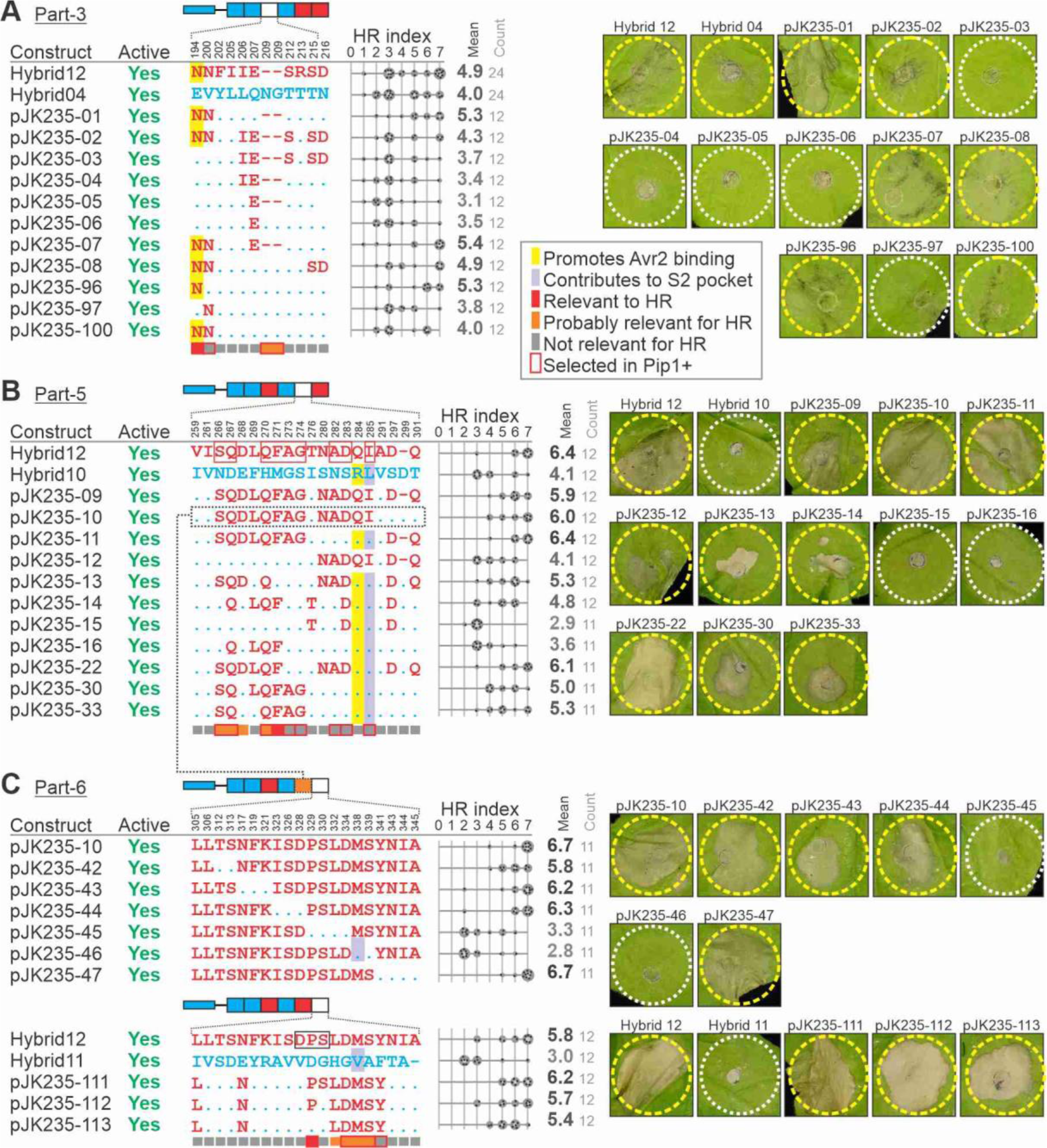
Further substitutions identify Rcr3 residues required for Avr2/Cf-2-dependent HR. Constructs tested to study variant residues in part-3 **(A),** part-5 **(B)** and part-6 **(C**) of the hybrid proteases. Shown on top is the general domain architecture of the listed constructs and below are the construct names; their activity detected by MV201 labelling; the variant residues in the respective parts with numbering of *Sl*Rcr3 on top; and the HR index upon co-expression with Avr2 and Cf-2 for n=12 replicates. Residues important for binding Avr2 are highlighted yellow and residues required for triggering Cf-2/Avr2-dependent HR, or possibly contribution to triggering Cf-2/Avr2-dependent HR are flagged with red and orange blocks under the alignments, respectively.

Part-5 contains 19 variant residues and one deletion in Rcr3. Given the large number of variants, we focused on residues predicted to be solvent-exposed in AlphaFold2-generated models of the Rcr3/Pip1 protein structures. Construct pJK235-11, carrying only the 10 solvent-exposed Rcr3-specific residues and the amino acid deletion, still triggers a full HR (**Figure 4B**; **Table S2**), suggesting that the remaining nine residues are not involved in triggering HR. However, constructs pJK235-09 and -10, which include five additional Rcr3-specific residues (NADQI), caused reduced HR activity compared to pJK235-11 (**Figure 4B**). This reduction correlates with the presence of Q284 from Rcr3, which is known to decrease Avr2 inhibition (Kourelis et al., 2020). Interestingly, constructs pJK235-09 and -10 display similar HR-inducing activity, indicating that the last three Rcr3-specific positions in part-5 (D-Q) do not significantly influence HR in the presence of Rcr3 residues NADQI. However, the comparison between constructs pJK235-11 with pJK235-30, suggests these residues (D-Q) do contribute quantitatively to HR (**Figure 4B**). Similarly, the comparison between constructs pJK235-12, which contains NADQI and D-Q, and pJK235-30, which includes only the first stretch (SQDLQFAG), indicates that the first stretch alone has a more pronounced effect on HR induction, whereas the latter two stretches contribute to a lesser, yet discernible extent when combined. Further analysis of constructs pJK235-14, pJK235-15, and pJK235-16 highlights the roles of individual residues. Construct pJK235-14, while weakly HR-active, differs significantly in HR induction from pJK235-15 and pJK235-16 (**Figure 4B**), underscoring the importance of Q267 and/or the LQF sequence. The absence of either S266 and/or A273 and G274 requires either D283 and/or D297 for effective HR induction (**Figure 4B**). Moreover, construct pJK235-22 triggers full HR (**Figure 4B**), confirming that A273 and G274 are non-essential for HR. Its higher HR activity compared to pJK235-13 implies that L269 and/or F271 are contributors. Yet, the comparison of constructs pJK235-33 and pJK235-30 reveals that L269 does not contribute significantly, indicating a role for F271 in HR induction in pJK235-22. In summary, while most of the 19 variant positions in part-5 do not contribute significantly to HR, residues S266, Q267, D268, Q270, and F271 (SQD-QF) collectively contribute to HR. Additionally, Pip1 residue R284 contributes to HR quantitatively by enhancing Avr2 inhibition, along with quantitative contributions to HR by N280/A282/D283, and D297/Q302.

Finally, part-6 contains 19 variant residues and one insert/deletion amino acid. Part-6 also contains the DPS insertion required for *Nicotiana* Rcr3s to trigger Avr2/Cf-2-dependent HR (**Figure 2**), but this sequence is VDG in Pip1. To further pinpoint residues in part-6 we first tested the effect of replacing six regions with Pip1 residues in pJK235-10, which also carries seven Pip1-specific residues in part-5. This scan revealed that Rcr3-specific residues TS, NFK, ISD and YNIA are not essential for HR, and that Rcr3-specific residues PSLD and MS contain residues that are essential for triggering HR (**Figure 4C**; **Table S3**). Thus, the D328 in the DPS motif is not required for triggering HR. Constructs pJK235-111 and -112 have similar HR-inducing activity (**Figure 4C**), indicating that S330 in the DPS motif is also not required. pJK235-113, however, is less able to trigger HR (**Figure 4C**), which indicates that P329 in the DPS motif is important for HR. In conclusion, P329 and the combination of four variant residues (LDMS) are the only four Rcr3-specific residues contributing to HR in part-6.

### Pip1+ differs by only 18 residues from Rcr3 and triggers Avr2/Cf-2-dependent HR

Whilst we were still analysing the role of the variant residues within parts-3, -5 and -6, we generated Pip1+, which is a Pip1 protease that contains only 18 Rcr3-specific residues (**Figure 5A**), including 11 of the 13 variant residues that were associated with HR. The two missing residues are D268 and L332, which were not tested separately but were found to contribute to HR in conjunction with other variant residues. The selection of these 18 residues involved systematically removing Rcr3-specific residues from parts -3, -5, and -6 through a three-step process: firstly, by omitting residues with less than 50% exposure in Rcr3 structural models; secondly, by excluding variants present in natural Rcr3 orthologs from *Solanum* species; and thirdly, by removing chemically similar variants in Pip1 orthologs (Supplemental **Figure S5**). Pip1+ therefore contains N194 (enhances Avr2 binding); N200; the two-residue deletion in part-3 (ΔNG); S266; Q267; Q270; F271; A273; G274; A282; D283; I285, P329, D334, M338, S339 and Y341. Pip1+ still has 78 (84%) of the variant residues in the protease domain, and shares 94.8% identity with Pip1 across the entire protein and only 53.8% with Rcr3. Importantly, the co-expression of Pip1+ with Avr2 and Cf-2 triggers HR with similar strength as Rcr3 (**Figure 5B**), confirming that most of the variant residues do not contribute to HR.

**Figure 5.**
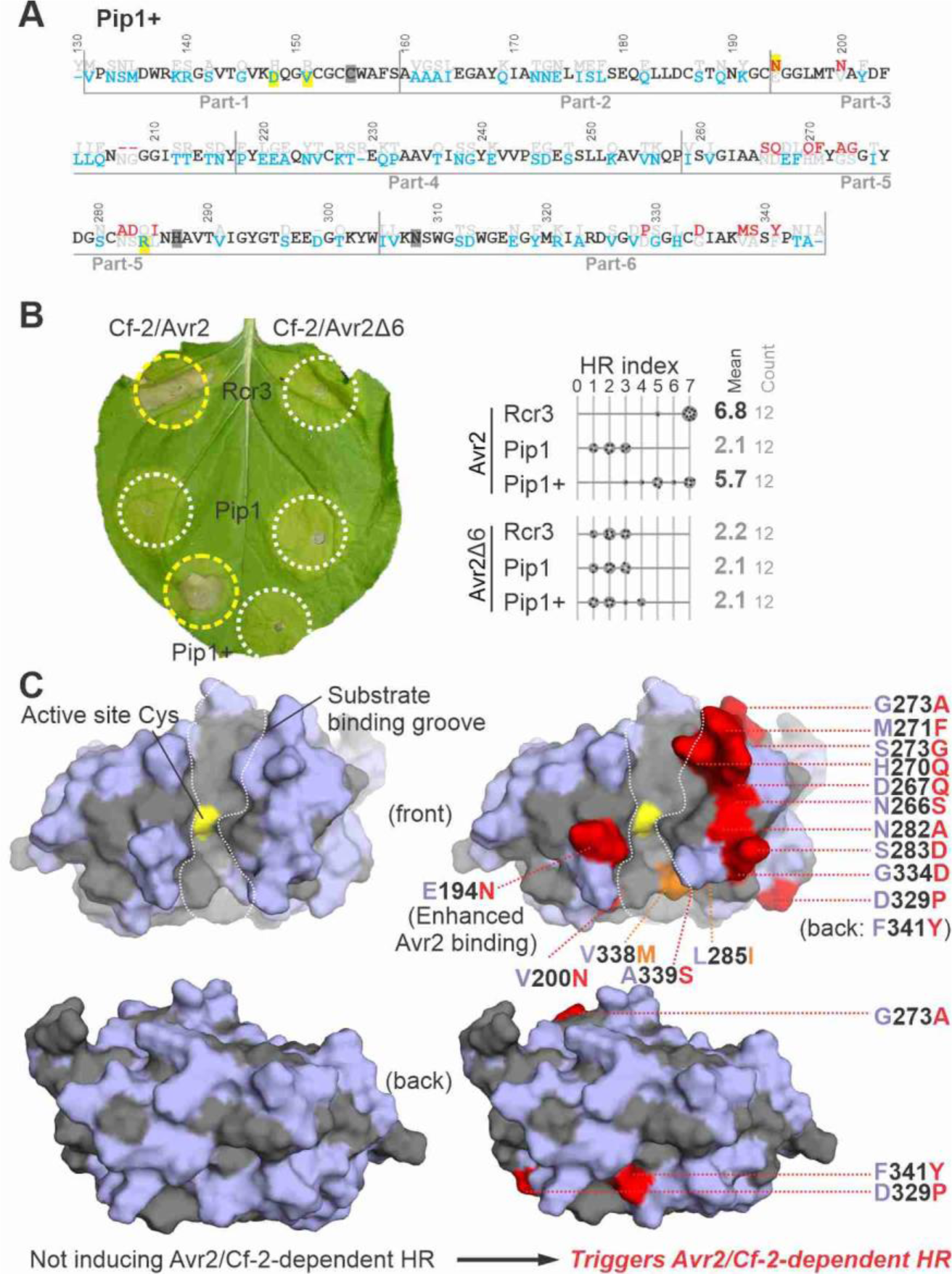
Pip1+ contains only 18 Rcr3-specific positions and triggers Avr2/Cf-2-dependent HR. **(A)** Protein sequence of the protease domain of Pip1+, showing residues specific to Rcr3 (red), Pip1 (blue) or both (black). The four residues promoting Avr2 affinity are highlighted in yellow and the active site residues in grey. **(B)** Pip1+ triggers Avr2/Cf-2-dependent HR. Rcr3, Pip1 and Pip1+ were transiently co-expressed with Cf-2 and Avr2/Avr2Δ6 in *N. benthamiana* by agroinfiltration. Shown is a representative leaf with infiltrated regions encircled with a white line (no HR) or yellow line (HR). The HR index was determined using n=12 replicates. **(C)** Structural models of Pip1 and bioengineered Pip1+. Both sequences were modelled using AlphaFold2 (both pTM= 0.94), and presented in PyMol using surface representation of the front of the protease with residues that are identical to tomato Rcr3 (grey) and that are specific to Pip1 (blue). Bioengineering of Pip1 with 16 Rcr3-specific residues (red/orange) causes Avr2/Cf-2-dependent HR. Two Rcr3-specific substitutions (V338M and L285I, orange) alter substrate specificity.

Modelling the structure of both Pip1 and the engineered Pip1+ with AlphaFold2 resulted in reliable complexes (pTM = 0.94), that show that the variant residues in Pip1 when compared to tomato Rcr3 scatter over the surface on both sides of the protein (**Figure 5C**, red residues). Most of these variant residues, however, are not required for Avr2/Cf-2-dependent HR. Pip1 carries VDG instead of the DPS insertion missing from *Nt*Rcr3 but a D329P substitution in this extended loop is essential to induce Avr2/Cf-2-dependent HR. It seems likely that the P329 proline residue will restrict the folding flexibility of the VDG loop but such structural change was not predicted using Alphafold2 modelling. Pip1 requires N194 to enhance Avr2 binding and trigger Avr2/Cf-2-dependent HR, similar to *Nt*Rcr3. The remaining residues required for Avr2/Cf-2-dependent HR in Pip1 all cluster on the top of the β-sheet lobe (**Figure 5C**), which represents a likely platform for interactions with Cf-2.

### Pip1 and Rcr3 have distinct substrate specificities

To investigate whether Rcr3 and Pip1 may have distinct substrate specificities, we used a modified version of proteomic identification of protease cleavage sites (PICS) with an *E. coli* proteome-derived peptide library generated by digestion with trypsin and Lys-C (Biniossek et al., 2016) (**Figure 6A**). With this library we assessed substrate specificity of recombinant Pip1, Rcr3 and *Sl*Rcr3, which were produced in *Pichia pastoris* and purified via a C-terminal His-tag (Paulus et al., 2020). As with other papain-like Cys proteases, Rcr3 and Pip1 also display selectivity for a hydrophobic P2 residue in the substrate (**Figure 6B**), but the selectivity is distinctly different. Both Rcr3 and *Sl*Rcr3 share a preference for P2 being either proline (P), methionine (M), valine (V) or leucine (L) (**Figure 6B**). By contrast, Pip1 prefers P2 being either leucine (L) or phenylalanine (F) (**Figure 6B**). This indicates that Rcr3 and Pip1 have distinct substrate specificities, and therefore likely have distinct substrates.

**Figure 6.**
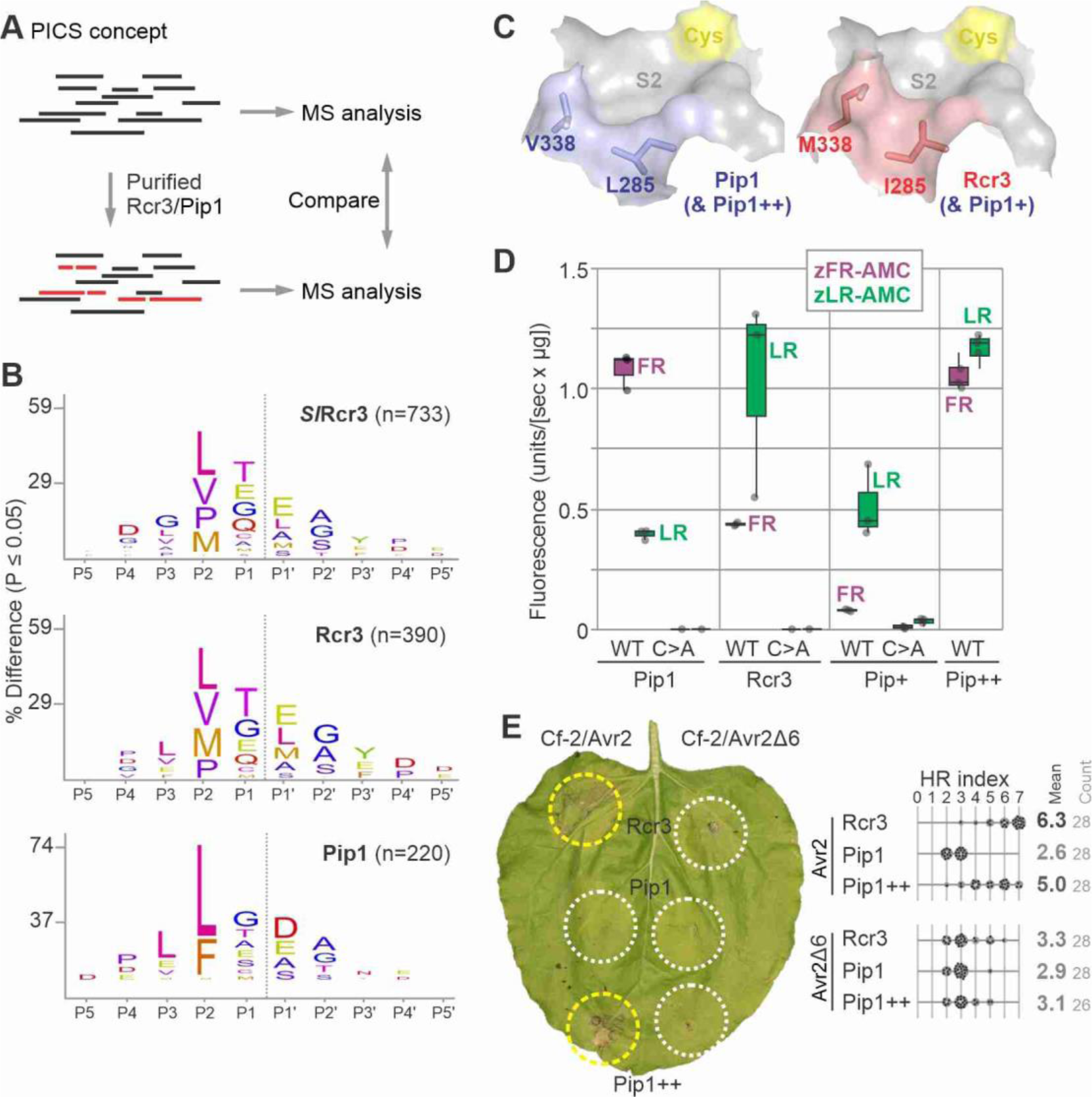
S2 pocket bioengineering reverts substrate specificity of Pip1+. **(A)** Determination of cleavage site preference using Proteomic Identification of Protease Cleavage Sites (PICS). A trypsin digest of *E. coli* is incubated with and without various proteases, isotopically labelled by reductive demethylation, mixed with the buffer control, and analysed by tandem MS. Differences with the internal buffer control are used to select cleaved peptides and generate the cleavage logo. **(B)** Cleavage logos for *Sl*Rcr3, Rcr3 and Pip1. A peptide library made using Trypsin/LysC from an *E. coli* proteome was digested with recombinant C-terminal 6xHis-tagged *Sl*Rcr3 and Pip1 purified from *Pichia pastoris*. This digested peptide library was di-methylated, mixed in equal amounts and analysed by LC-MS/MS. A custom PERL script was used to filter semi-specific peptides and through database-lookup determine the corresponding prime and nonprime sequence. Shown are the residues relative to the cleavable bond and their frequencies are used to scale the font size. The number of all and unique cleavage events that was used to calculate the cleavage logo are indicated. **(C)** Model for the different S2 pockets in Rcr3 and Pip1. Structural models were generated for Rcr3 and Pip1 with Alphafold2, and the S2 substrate binding pocket was visualised in PyMol with 50% surface presentation. Shown are the catalytic Cys (yellow), and the variant residues in Rcr3 (red) and Pip1 (blue). **(D)** Different dipeptide preferences for Pip1, Rcr3, Pip1+ and Pip1++. Pip1, Rcr3, Pip1+ and Pip1++ and their catalytic mutants were produced by agroinfiltration and purified via their C-terminal His tag. Equal protein amounts were incubated with zLR-AMC and zFR-AMC and fluorescence was read using a plate reader. The fluorescence intensity measured over 60 minutes was plotted for all samples. This experiment was repeated with independently produced proteases with similar results **(E)** Pip1++ triggers Avr2/Cf-2-dependent HR. Rcr3, Pip1 and Pip1++ were transiently co-expressed with Cf-2 and Avr2/Avr2Δ6 in *N. benthamiana* by agroinfiltration. Shown is a representative leaf with infiltrated regions encircled with a white line (no HR) or yellow line (HR). The HR index was determined using n=28 replicates.

**Figure 7.**
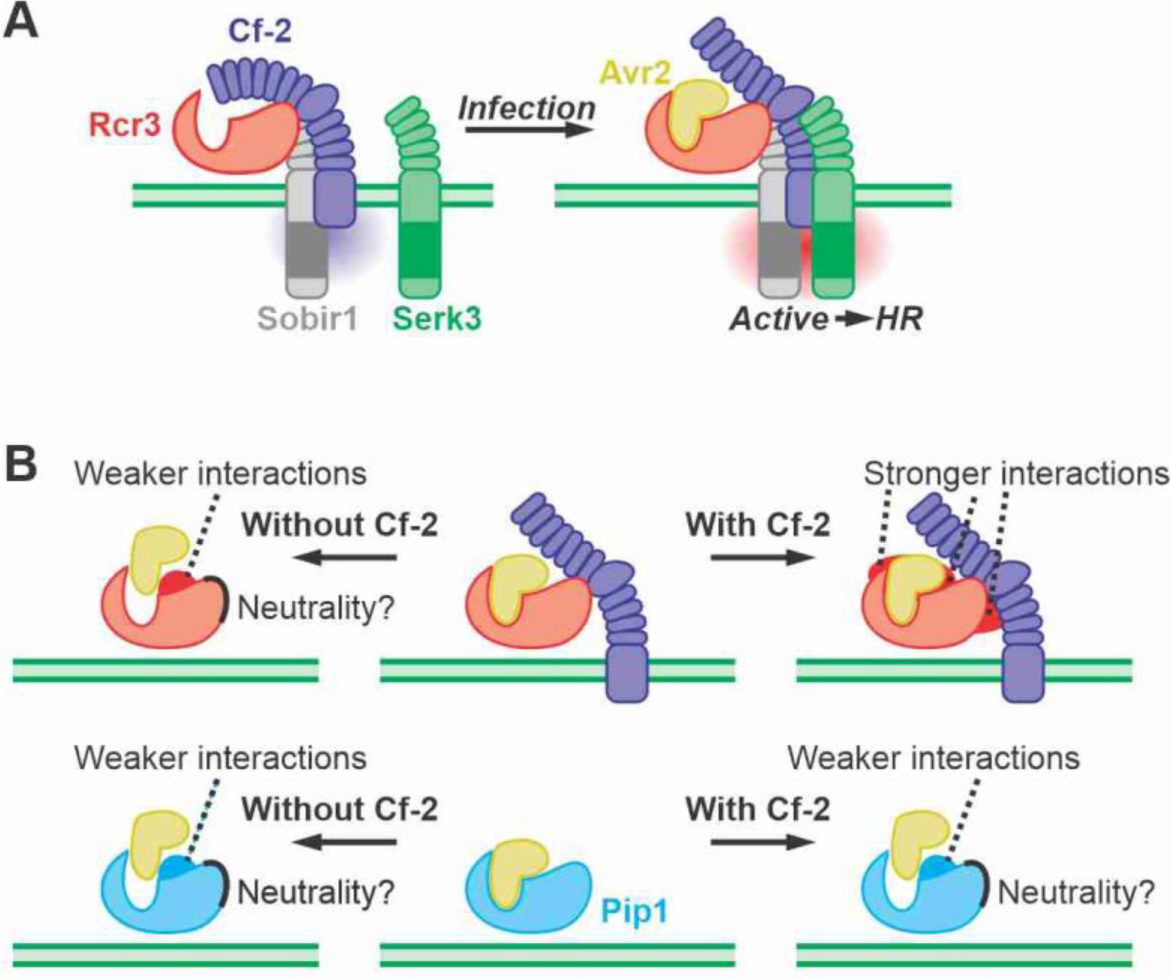
Working model of Avr2 perception and Rcr3 evolution. **(A)** In non-infected plants, Rcr3, Cf-2 and Sobir1 are interacting in a constitutive complex lacking Serk3. Upon infection, Avr2 would bind Rcr3 with high affinity, triggering a conformational change in Cf-2 that triggers the recruitment of Serk3, which then activates the hypersensitive response (HR). **(B)** Rcr3 is under opposing selection pressures in the presence/absence of Cf-2, at the putative interfaces with both Avr2 and Cf-2. By contrast, Pip1 will be under selection to avoid Avr2 inhibition and will probably not be under selection to interact with Cf-2.

We next analysed the structural models of Rcr3 and Pip1 to investigate the underlying reason for their differing substrate specificity. Variant residues I285L and M338V constitute the edge of the S2 pocket, opposite to the catalytic Cys (**Figure 6C**). The I285 and M338 residues in Rcr3 seem to keep the S2 pocket small, explaining the accommodation of smaller Val and Pro residues at the P2 position in substrates. By contrast, the L285 and V338 residues in Pip1 seem to extend the S2 pocket, explaining the accommodation of the much larger Phe(F) residues at the P2 position of the substrate. Thus, the putative larger S2 pocket in Pip1 might be why it can accommodate substrates with P2=Phe, in contrast to Rcr3, which prefers smaller residues at P2.

### Dipeptide substrates show that Pip1+ has an Rcr3 substrate preference

To confirm the difference in substrate specificity of Rcr3 and Pip1, we took advantage of two commercially available dipeptide substrates, z-FR-AMC and z-LR-AMC, which release fluorescent 7-amido-4-methylcoumarin (AMC) upon cleavage. According to PICS data, Rcr3 should preferentially cleave z-LR-AMC, whereas Pip1 should prefer z-FR-AMC. We transiently expressed Pip1 and Rcr3 by agroinfiltration and purified them using C-terminal His tags (Schuster et al., 2022). Catalytic mutants of each of these proteases were included as negative controls. Incubation of Rcr3 and Pip1 with these two substrates indeed confirmed their distinct substrate specificity: Rcr3 preferentially cleave z-LR-AMC and Pip1 preferentially cleaves z-FR-AMC (**Figure 6D**).

To test if Pip1+ has maintained the substrate specificity of Pip1, we produced Pip1+ and its catalytic mutant by agroinfiltration and purified it using a C-terminal His tag. Remarkably, enzymatic assays showed that Pip1+ preferentially cleaves z-LR-AMC, unlike Pip1 and similar to Rcr3 (**Figure 6E**). Thus by introducing Rcr3-specific residues into Pip1, we also changed the substrate specificity of Pip1.

### S2 pocket bioengineering reverts the substrate specificity of Pip1++

To test if the modified S2 pockets in Pip1+ are responsible for Rcr3-like substrate specificity, we generated Pip1++, which contains the L285 and V338 of Pip1 and is otherwise identical to Pip1+, carrying 16 Rcr3-specific residues. Enzymatic assays using purified proteases showed that Pip1++ exhibits significantly increased activity towards z-FR-AMC, consistent with the enlarged S2 pocket while retaining a high activity for z-LR-AMC (**Figure 6D**). This indicates that the S2 pocket can be bioengineered to revert the altered substrate specificity towards that of Pip1.

To determine if Pip1++ is also able to trigger HR with Avr2 and Cf-2, we co-expressed Pip1++ with Avr2 and Cf-2 by agroinfiltration and monitored the strength of HR. Pip1++ triggers a strong HR with Avr2 but not with Avr2Δ6, similar to Rcr3 (**Figure 6E**). Thus, although bioengineering the S2 pocket in Pip1+ changed its substrate specificity, it did not diminish its ability to trigger HR with Avr2 and Cf-2.

## DISCUSSION

We found that bioengineering eggplant Rcr3 with a single D244P substitution is sufficient to trigger Avr2/Cf-2-dependent HR. This residue does not affect the interaction with Avr2 and is likely to affect interaction with Cf-2. We also discovered that the insertion of three residues (D328/P329/S330) into *N. tabacum* Rcr3 or the resurrected *N. benthamiana* Rcr3, together with the G194N or K194N substitutions, are sufficient for Avr2/Cf-2-dependent HR. While the G194N and K194N substitutions are required for increased Avr2 binding, the D328/P329/S330 insertion are only required for HR and are absent from all *Nicotiana* Rcr3 homologs. In addition, a series of Rcr3/Pip1 hybrids revealed three regions in Rcr3 important for Avr2/Cf-2-dependent HR. Further fine-mapping uncovered that the HR-relevant residues cluster on one lobe of Rcr3, flanking the substrate-binding groove. Substitutions at the rim of the S2 pocket altered substrate specificity in Pip1+ but these substitutions are not essential for HR and reconstitute Pip1-like activity when mutated to Pip1-like residues in Pip1++.

Our findings lead us to propose a working model in which the Avr2/Rcr3 complex interacts with Cf-2 (**Figure 8A**). In the Pip1++ mutant, the residues essential for Avr2/Cf-2-dependent HR are predominantly located on the top of the β-sheet lobe next to the substrate-binding groove. This specific localization suggests that Cf-2 likely binds this region, enabling it to simultaneously detect the presence of Avr2 bound to the substrate-binding pocket of Rcr3. We hypothesize that the binding of the Avr2/Rcr3 complex induces a conformational change in Cf-2, triggering HR. This process likely involves recruiting SERK3 (also known as BAK1), as observed in other receptor-like proteins, including Cf-4 (Postma et al., 2016). This activated immune complex would lead to the activation of NRC3, thereby inducing HR and immunity towards the pathogen (Kourelis et al., 2022).

The presence of Rcr3 and Pip1 in solanaceous plants whilst Cf-2 is specific to the *Solanum* genus (Kourelis et al., 2020) indicates that Rcr3 is under diverse selection pressures in different plant species. In *Solanum* plants that have Cf-2, selection pressure will be on increasing Avr2/Rcr3 and Rcr3/Cf-2 interactions, whilst in the absence of Cf-2 in *Solanum* plants, the selection pressure will be on Rcr3 to avoid inhibition by Avr2 (**Figure 8B**). This should cause opposing selection forces in two distinct areas of Rcr3: residues required for Cf-2-dependent HR will be under selection to increase interactions with Cf-2 only in *Solanum* and evolve under neutrality in other solanaceous plants. This could explains why eggplant and tobacco Rcr3 carry D244 and the DPS deletion, respectively, which both abolish HR signalling with Cf-2. Further evolutionary studies of Rcr3 in wild solanaceous plants may elucidate the ancestral state of Rcr3 and may show how Rcr3 is adapted in *Solanum* to mediate Cf-2-dependent HR signalling. The second area under selection in Rcr3 are the residues around the substrate binding groove where inhibitors bind: the ‘ring-of-fire’ (Hörger and van der Hoorn, 2013). These ‘ring-of-fire’ positions are indeed highly variant (Shabab et al., 2008) and under diversifying selection in Rcr3 (Hörger et al., 2012). Indeed, four of these variant residues affect Avr2 affinity (Kourelis et al., 2020). These, and other variant residues might also affect interactions with other Rcr3 inhibitors such as Cip1 and EpiCs (Song et al., 2009; Shindo et al., 2016; Schuster et al., 2024). Further studies of Rcr3 in wild solanaceous plants could reveal the different selection pressures on these residues, depending on the presence of Cf-2 and the inhibitors used by different pathogens. Although the selection pressure on Pip1 is much less than on Rcr3, variant residues also accumulated around the substrate binding groove (Shabab et al., 2008), consistent with arms races with various pathogen-derived inhibitors. Modification of two of these residues (D147R, V150E) renders Pip1 insensitive to Pip1 inhibitor EpiC2B (Schuster et al., 2024). Residues that are required for Cf-2-mediated HR in Rcr3 might be under neutrality or negative selection in Pip1, although it is interesting that the only variant residue between Pip1 and *Sp*Pip1 (H329D) is at one such position. Further studies on the occurrence of variant residues in Rcr3 and Pip1 and their relevance to disease and immunity may uncover strengths and directionalities of selection on these immune proteases.

Our attempts to physically demonstrate interactions of Avr2/Rcr3 or Rcr3 with Cf-2, including co-immunoprecipitation assays and blue native gel electrophoresis, have been unsuccessful so far, despite the availability of active Avr2 and Rcr3 proteins (e.g. Paulus et al., 2020), and a good expression of the soluble ectodomain of Cf-2 in *N. benthamiana*. This suggests that additional specific conditions or cofactors might be required to recapitulate the Avr2/Rcr3/Cf-2 complex in *in vitro* assays. Although the interactions between some pathogen-secreted effector proteins and proteases such as the tomato-secreted P69B protease can be successfully predicted with AlphaFold Multimer (AFM; Homma et al., 2023), we were unsuccessful in similarly predicting the structure of the Avr2/Rcr3/Cf-2 complex. The inability of AFM to predict a reliable structure of Avr2 is probably caused by the low number of available Avr2-homologous sequences, whilst the large leucine-rich repeat (LRR) region of Cf-2 can easily support false negative complexes predicted by AFM. Interactions between Cf-2 and Avr2/Rcr3 are likely but remain to be demonstrated.

*Sl*Rcr3 is able to trigger Cf-2-dependent auto-necrosis in tomato (Krüger et al., 2002) but this can be suppressed in hybrids that co-express the Rcr3 allele, which co-evolved with Cf-2 in *S. pimpinellifolium* (Ilyas et al., 2015). This indicates that *Sl*Rcr3 has a lower affinity for Cf-2 and can be outcompeted by Rcr3. The ability of *Sl*Rcr3 to trigger auto-necrosis with Cf-2 is only detected in tomato and could not be established in *N. benthamiana* (Kourelis et al., 2020), which might suggest that tomato expresses a co-factor that binds *Sl*Rcr3 more strongly than Rcr3 and triggers HR in a similar way as Avr2 does. Rcr3 has also been reported to trigger Cf-2-dependent HR with potato cyst nematode (*Globodera rostochiensis*) effector *Gr*VAP1 (Lozano-Torres et al., 2012), but we were unable to reproduce these results (Kourelis et al., 2020). By contrast, both the *Pseudomonas syringae* pv. *tomato* DC3000 chagasin-like effector Cip1, and the *Phytophthora infestans* cystatin-like effectors EpiC1 and EpiC2B can inhibit Rcr3, but they do not trigger *Cf-2*-dependent HR (Shindo et al., 2016; Song et al., 2009). One reason for not triggering Cf-2-dependent HR might be that Cf-2 may not be able to directly interact with the effector in the Cip1/Rcr3 and EpiC/Rcr3 complexes. A second reason might be that both Cip1 and EpiCs are relatively weak inhibitors of Rcr3, as they both preferentially inhibit immune protease C14 (Kaschani et al., 2010; Shindo et al., 2016). Avr2 has a much higher affinity for Rcr3 (Kaschani et al., 2010; Shindo et al., 2016), and might be able to insert itself into a preformed Rcr3/Cf-2 complex, causing a conformational change in Cf-2 that triggers HR. The latter model would be consistent with the presence of an endogenous inhibitor of tomato that binds stronger to *Sl*Rcr3 than to Rcr3, explaining the auto-necrosis induced by *Sl*Rcr3 but not Rcr3. This model indicates that bioengineered Rcr3 with increased affinity to Cip1 and/or EpiCs could confer Cf-2-dependent immunity to *Pseudomonas syringae* and/or *Phytophthora infestans*, respectively. Although the Rcr3/Pip1 hybrids generated in this study were not able to induce Cf-2-dependent HR in the presence of either Cip1 or EpiCs (Supplemental **Figure S4**), we did not yet generate and test Rcr3 variants with an increased affinity to these effectors.

A similar concept of ‘decoy bioengineering’ has previously been demonstrated for pseudokinase PBS1, which is a decoy in the recognition of AvrPphB, a type-III protease effector produced by *Pseudomonas syringae*, by resistance protein RPS5, a Nod-like Receptor (NLR) in Arabidopsis (Kim et al., 2016). PBS1 contains a cleavage site for AvrPphB and replacing this cleavage site with that of other pathogen-produced proteases triggered RPS5 activation by proteases of these other pathogens, resulting in immunity to *P. syringae* expressing the AvrRpt2 protease effector, or to turnip mosaic virus (TuMV) expressing its viral protease (Kim et al., 2016; Pottinger et al., 2020). Interestingly, in other plant species, PBS1 homologs are similarly guarded by unknown NLRs through a process of convergent evolution (Carter et al., 2019). Bioengineering these PBS1 homologs in soybean or potato can also provide immunity against different viral pathogens, even without knowing the identity of the NLR guarding these PBS1 homologs (Helm et al., 2019; Pottinger et al., 2020; Bai et al., 2022). This suggests that it is possible that decoy bioengineering could provide an alternative route to an altered recognition spectrum.

Indeed, some NLRs are able to interact with different decoys in nature. One notable example is the highly conserved NLR ZAR1 (Adachi et al., 2023; Gong et al., 2022), which interacts with different (pseudo)kinases to recognise different effectors (Lewis et al., 2013; Wang et al., 2015; Seto et al., 2017; Schultink et al., 2019; Laflamme et al., 2020; Martel et al., 2020). In the case of ZAR1, the (pseudo)kinases appear to be under evolutionary pressure to diversify (Gong et al., 2022). Similarly, Rcr3 is under selection to diversify at residues surrounding the substrate binding groove (Hörger et al., 2012), possibly to create alleles that confer recognition of other pathogen-secreted protease inhibitors. Decoy bioengineering may provide an alternative route for the generation of synthetic resistance genes targeting important plant pathogens in the future. Finally, it is important to note that Cf-2 is not the only cell-surface receptor that has been shown to require additional apoplastic factors for ligand recognition, as the apoplastic protein auxin-binding protein 1 (ABP1) and ABP1-LIKE 1 and 2 (ABL1/2) physically interacts with the LRR-RLK transmembrane kinase (TMK) family receptor-like kinases to form an auxin-sensing complex in the apoplast (Yu et al., 2023).

Our enhanced understanding of Rcr3 can now be applied to exploit the Rcr3/Cf-2 perception system to provide immunity to different pathogens through decoy bioengineering. PLCPs, such as Pip1 and Rcr3 are universally secreted by plants, including in key crops like maize, citrus, rice, and papaya where they play a vital role in defending the apoplast against pathogen colonization (Misas-Villamil et al., 2016). In response, many host-adapted apoplastic pathogens secrete PLCP inhibitors to facilitate colonization, as seen in pathogens such as the maize smut pathogen *Ustilago maydis* (Mueller et al., 2013), the citrus Huanglongbing pathogen *Candidatus Liberibacter asiaticus* (Clark et al., 2018), the crucifer clubroot pathogen *Plasmodiophora brassicae* (Pérez-López et al., 2021), the tomato bacterial spot pathogen *Pseudomonas syringae* (Shindo et al., 2016), and the potato late blight pathogen *Phytophthora infestans* (Tian et al., 2007). Bioengineering the Rcr3/Cf-2 perception system to recognise these PLCP inhibitors can be an effective and sustainable approach to develop a perception system that confers recognition of the many different bacterial, fungal and oomycete pathogens that secrete PLCP inhibitors when colonising the apoplast.

## MATERIAL & METHODS

### Plant material and growth conditions

*N. benthamiana* LAB and *S. lycopersicum* MoneyMaker (MM)-*Cf-2 rcr3-3* (Dixon et al., 2000) plants were propagated in a glasshouse and were grown in a controlled growth chamber with temperature 22–25°C, humidity 45–65% and 16/8 hr light/dark cycle.

### General plasmid constructions

The Golden Gate Modular Cloning (MoClo) kit (Weber et al., 2011) and the MoClo plant parts kit (Engler et al., 2014) were used for cloning, and all vectors are from this kit unless specified otherwise. Unless stated otherwise, receptors, effectors, and Rcr3 variants were cloned into the binary vector pJK268c, which contains the tomato bushy stunt virus silencing inhibitor p19 in the backbone (Paulus et al., 2020). Cloning design and sequence analysis were done using Geneious Prime (v2022.2.1; https://www.geneious.com). Plasmid construction is described in Supplemental **Table S4**.

### Fragment-swapped hybrid cloning

For fragment-swapping the nucleotide region encoding for the Rcr3 and Pip1 peptidase C1 domain were divided into six arbitrary fragments and ordered through gene-synthesis (Biomatik; pJK103-pJK107, pJK111-pJK115, pJK185, pJK186; Supplemental **Table S4**). These fragments, together with a fragment containing the nucleotides encoding for the Pip1 pro-domain but lacking the signal peptide (pJK110; Supplemental **Table S4**) were combined in all 64 possible combinations with pICH41264 (Addgene #4799; Weber et al., 2011) in a BpiI Golden Gate reaction (Supplemental **Table S4**). These fragment-swapped level 0 modules were subsequently combined with pJK001 and pJK002 (Grosse-Holz et al., 2018), and pICH51288 and pICH41414 (2x35S and 35S terminator; Addgene #50269 and #50337; Engler et al., 2014) in a BsaI Golden Gate reaction to generate binary vectors containing the fragment-swapped Rcr3/Pip1 hybrids driven by the double 35S CaMV promoter and targeted to the apoplast using a *Nt*PR1a signal peptide. All used and created plasmids are summarized in Supplemental **Table S4**.

### *E. coli* expression vector cloning Avr2, Cip1, EpiC1, and EpiC2B purification

The expression vectors for periplasmic secretion of N-terminal 6xHis-tagged Avr2 (pJK153; pET28b-T7::OmpA-6xHis-TEV-Avr2) or Avr2Δ6 (pSM101; pET28b-T7::OmpA-HIS-TEV-Avr2Δ6) (Kourelis et al., 2020), and Cip1 (pJK159; pET28b-T7::OmpA-6xHis-TEV-Cip1), or C-terminal 6xHis-tagged EpiC1 (pJK254; pET28b-T7::OmpA-EPIC1-6xHis) and EpiC2B (pJK256; pET28b-T7::OmpA-EPIC2B-6xHis), were generated as described in Supplemental **Table S4**, and transformed into *E. coli* strain Rossetta2(DE3)pLysS (Novagen/Merck).

For protein purification, an overnight grown starter culture was diluted 1/100 in TB [per liter: 24 g yeast extract, 12 g bacto-tryptone, 0.5% glycerol, after autoclaving, 0.2 M/L potassium phosphate buffer [26.8 g KH_2_PO_4_ and 173.2 g K_2_HPO_4_ pH 7.6] was added and the bacteria were grown for ∼3 h to OD_600_ ≈ 0.6 and induced overnight at 37 °C upon adding 0.8 mM isopropyl β-D-1-thiogalactopyranoside (IPTG). The supernatant was collected by centrifuging the bacterial culture at 10,000 × *g*, 4 °C for 20 min, and was adjusted to 50 mM Tris-HCl, 150 mM NaCl, 10 mM imidazole and run over a Ni-NTA agarose (Qiagen)-loaded column by gravity-flow. The Ni-NTA agarose was washed with 10 column volumes washing buffer (50 mM Tris-HCl, 150 mM NaCl, 50 mM imidazole, pH 8) and the protein was eluted using elution buffer (50 mM Tris-HCl, 150 mM NaCl, 250 mM imidazole, pH 8). The purified protein was concentrated by centrifugation in 3 kDa cut-off filters (Amicon Ultra-15 Centrifugal Filter Units, Merck) at 4500 × *g*, 4 °C, flash-frozen in liquid nitrogen, and stored at −80 °C. Protein concentration was determined by Bradford assay using BSA as a standard, and purity was verified by running the protein on 18% SDS-PAGE gels followed by Coomassie staining.

### Transient gene-expression and cell death assays

Transient gene expression in *N. benthamiana* was performed by agroinfiltration according to methods described by (van der Hoorn et al., 2000). Briefly, *A. tumefaciens* strain GV3101 pMP90 carrying binary vectors were inoculated from glycerol stock in LB supplemented with appropriate antibiotics and grown overnight at 28 °C until saturation. Cells were harvested by centrifugation at 2000 × *g* at room temperature for 5 min and resuspended in infiltration buffer (10 mM MgCl_2_, 10 mM MES-KOH pH 5.6, 200 µM acetosyringone) to OD_600_ 0.25 for each construct in the stated combinations and left to incubate in the dark for 2 h at room temperature prior to infiltration into young, fully expanded leaves of 5-week-old *N. benthamiana* plants. Replicate numbers given indicate independent infiltration of different leaves from multiple plants. For quantitation, HR cell death phenotypes were scored in a range from 0 (no visible necrosis) to 7 (fully confluent necrosis) as in Kourelis et al., 2022 and plotted using a R script modified from Bentham et al., 2023. Statistical analysis was conducted using the besthr R package (MacLean, 2020).

### HR assays in tomato

Apoplastic fluid (AF) was isolated from agroinfiltrated *N. benthamiana* plants expressing different Rcr3/Pip1 protein variants at 5 days post-infiltration. Briefly, the AF was extracted by vacuum infiltrating *N. benthamiana* leaves with ice-cold MilliQ. Leaves were dried with filter paper to remove excess liquid, and apoplastic fluid was extracted by centrifugation of the leaves in a 20 ml syringe barrel (without needle or plunger) located in a 50 ml falcon tube at 2000 × *g*, 4 °C for 25 min. 1 µM Avr2 (final concentration) was added to this AF, which was infiltrated into leaflets of tomato MM-*Cf-2 rcr3-3* plants (Dixon et al., 2000). Leaves were imaged 2-5 dpi.

### Activity-based protein-profiling

AF was isolated from agroinfiltrated *N. benthamiana* plants expressing different proteins at 2–3 days post-infiltration and samples were used directly in labelling reactions. Samples were adjusted to 5 mM DTT, 50 mM NaAc pH 5.0, and preincubated for 45 min with either 100 µM E-64 (a general PLCP inhibitor), 2 µM purified Avr2 or Avr2Δ6, or 1% DMSO (final concentration) as a control. After preincubation, samples were labelled with 2 µM MV201 for 3 h at room temperature (total volume 50 µl) (Richau et al., 2012). The labelling reaction was stopped by precipitation with 5 volumes of ice-cold acetone, followed by a 10 sec vortex and immediate centrifugation at 16,100 × *g*, 4 °C for 5 min. Samples were resuspended in a 2× sample buffer [100 mM Tris-HCl (pH 6.8), 200 mM DTT, 4% SDS, 0.02% bromophenol blue, 20% glycerol] and boiled for 5 min 95 °C prior to separation on 15% SDS-PAGE gels. Fluorescence scanning was performed on an Amersham Typhoon-5 scanner (GE Healthcare), using Cy3 settings. Equal loading was verified by Coomassie staining (Pink et al., 2010).

### *P. pastoris* expression vector cloning and protein purification

The *Pichia pastoris* PichiaPink™ yeast expression system was used to produce C-terminally 6xHis-tagged *Sl*Rcr3 (pPINKα-HC-*Sl*Rcr3-6xHis), Rcr3 (pJK206; pPINKα-HC-Rcr3-6xHis), and Rcr3(C153A/C154A) (pJK209; pPINKα-HC-Rcr3(C153A/C154A)-6xHis) (Paulus et al., 2020), or Pip1 (pJK211; pPINKα-HC-Pip1-6xHis) (Supplemental **Table S4**). The proteases were cloned after the dibasic KR Kex2 protease cleavage site of the N-terminal secretion signal from the *S. cerevisiae* α-factor, and were driven by the methanol-induced *AOX1* promoter. 10 µg of plasmid DNA linearized using SpeI was transformed into electrocompetent PichiaPink™ strain 4 (ThermoFisher; *P. pastoris* genotype *ade2*, *pep4*, *prb1*) targeted to the *TRP2* locus, followed by selection on PAD selection plates (ThermoFisher) and selection of white colonies indicative of multiple integrations. Glycerol stocks were prepared by concentrating the transformants to a final OD_600_ of 50–100 (approximately 2.5–5.0 × 10^9^ cells/ml) in 25% glycerol and aliquots were snap frozen in liquid nitrogen and stored at -80°C.

For protein expression, a single glycerol stock was thawed and used to inoculate a 25 ml starter culture of BMGY medium (1% yeast extract; 2% peptone; 100 mM potassium phosphate, pH 6.0; 1.34% yeast nitrogen base (YNB) with ammonium sulphate and without amino acids; 0.00004% biotin; 1% glycerol) in a 250 ml baffled flask, grown at 30°C, 300 rpm until the culture reached an OD_600_ of 2–6 (approximately 2–3 days). This starter culture was used to inoculate 1 l of BMGY divided over four 1 l baffled flasks and grown at 30°C, 300 rpm until the culture reached log phase growth (OD_600_ = 2–6). Cells were harvested by centrifugation in sterile centrifuge bottles at 3,000 x g for 5 min at room temperature. To induce expression, the supernatant was decanted, and the pellet was resuspended in 200 ml of BMMY medium (1% yeast extract; 2% peptone; 100 mM potassium phosphate, pH 6.0; 1.34% yeast nitrogen base (YNB) with ammonium sulphate and without amino acids; 0.00004% biotin; 0.5% methanol). This culture was divided between two 1 l baffled flasks which were covered with two layers of sterile gauze, and grown for 24 hours at 30°C, 300 rpm. Afterwards the supernatant was collected by centrifugation of the culture at 10,000 x g, 4°C for 20 min. The supernatant was adjusted to 150 mM NaCl, 30 mM imidazole and run over a Ni-NTA agarose (Qiagen)-loaded column by gravity-flow. The Ni-NTA agarose was washed with 10 column volumes of washing buffer (50 mM Tris-HCl, 150 mM NaCl, 50 mM imidazole, pH 8) and the protein was eluted using elution buffer (50 mM Tris-HCl, 150 mM NaCl, 250 mM imidazole, pH8). Finally, the purified protein was concentrated by centrifugation in 3 kDa cut-off filters (Amicon Ultra-15 Centrifugal Filter Units, Merck) at 4,500 x g, 4°C, flash-frozen in liquid nitrogen and stored at -80°C. Protein concentration was determined by Bradford assay using BSA as a standard, and purity was verified by running the protein on 15% SDS-PAGE gels followed by Coomassie staining.

### Proteomics Identification of Cleavage Sites (PICS) peptide library preparation

Proteome- derived peptide libraries were produced essentially as described (Demir et al, 2022). Briefly, proteins were extracted from an E.coli cell pellet in 6 M Guanidine-HCl,100 mM HEPES, 150 mM NaCl, adjusted to pH 7.5 and supplemented with 10 mM EDTA and 1 mM PMSF directly before use. Proteins were reduced with 10 mM dithiothreitol (DTT) and alkylated with 40 mM iodoacetamide at 25°C, the reaction quenched after 60 min with another 10 mM DTT. Proteins were purified by chloroform/MeOH precipitation, resuspended in 100 mM HEPES pH 7.5 and digested at 37°C overnight with Trypsin/Lys-C (Promega) at a protease-to-proteome ratio of 1:100 (wt/wt). A small aliquot was analysed by SDS-PAGE to assure complete digestion. Digestion proteases were inhibited with 1 mM PMSF and heating to 70°C for 20 min, the sample acidified with 0.5% (vol/vol) TFA and degassed by application of a mild vacuum. Peptides were purified on Sep-Pak C18 solid-phase extraction cartridges (Waters) according to the manufactures protocol and the resulting peptide library stored in aliquots at a peptide concentration of 2 mg/ml at -80°C.

### PICS specificity assay

100 μg of the PICS peptide library was used per treatment. The peptide library was diluted to 1 mg/ml (final concentration) in 50 mM NaAc pH 5 and 10 mM tris(2-carboxyethyl)phosphine (TCEP; final concentration). Purified protease (Rcr3-His, *Sl*Rcr3-His or Pip1-His) or control (buffer) were matured in 50 mM NaAc pH 5 and 10 mM TCEP (final concentration) for 5 hours, prior to addition at a ratio of 1:100 (wt/wt) and incubation for 1hr at room temperature. 100 μM E-64 (final concentration) was added to inactivate the proteases, and proteases were further heat-inactivated for 5 min at 95°C. Protease-treated and control samples were isotopically labelled by reductive dimethylation. Rcr3-, *Sl*Rcr3-, Pip1-, and buffer-treated samples were labelled with either 20 mM ^12^CH_2_O (cat. no. 252549, Sigma-Aldrich) and 20 mM NaBH_3_CN (cat. no. 252549, Sigma-Aldrich) (light), 20 mM ^12^CD_2_O (Sigma-Aldrich) and 20 mM NaBH_3_CN (medium), or 20 mM ^13^CD_2_O (cat. no. 596388, Sigma-Aldrich) and 20 mM NaBD_3_CN (Sigma-Aldrich) (heavy), respectively, and incubated overnight at 37°C. Afterwards an additional 20 mM formaldehyde (final concentration 40 mM) and 20 mM sodium cyanoborohydride (40 mM final concentration) was added to ensure complete labelling. Labelling was quenched by the addition of 100 mM Tris-HCl pH 6.8 and 60 min incubation at 37°C. Following labelling, protease-treated and control samples were mixed at equal ratios and desalted by C18 solid phase extraction (Sep-Pak, Waters) according to manufacturer’s instructions and concentrated to 10 μl using a vacuum concentrator (ThermoFisher).

### Mass spectrometry data acquisition and analysis

For MS analysis of the PICS samples, 3 μl of the desalted samples were loaded onto a C18 reverse phase capillary trap column (Acclaim PepMap C18, 75 μm x 2 cm, 3 μm particle size, Thermo) and separated using a C18 reverse phase analytical column (Acclaim PepMap C18, 75 μm x 25 cm, 2 μm particle size, 100 Å, Thermo) coupled to an UltiMate3000 nano RSLC system (Thermo). A 90 min gradient from 5 to 32% (v/v) acetonitrile 0.1% formic acid in H_2_O was used to elute the peptides at a flow rate of 300 nl min^-1^. The nano-LC system was on-line coupled to an Impact II high-resolution quadrupole-time of flight tandem mass spectrometer (Bruker) using a CaptiveSpray nano-electrospray source (Beck et al., 2015). MS spectra were acquired in a range from m/z 200 to 1750 at 4 Hz. For fragmentation, the 17 most intense precursor ions of each MS scan were selected (Top 17 method). MS/MS spectra of the fragmented precursor ions were recorded in a mass range from m/z 300 to 1750 at an intensity-dependent collection rate of 2 to 20 Hz spectrum. The sample was injected twice.

Peptides were identified and quantified using the MaxQuant software package, version 1.6.0.16 (Tyanova et al., 2016). Generic settings for Bruker Q-TOF instruments were used to match spectra to protein sequences to the *E. coli* proteome database (strain K12, reference proteome, 4308 entries). Search parameters included precursor mass tolerance of ±10 ppm, fragment ion mass tolerance of ±20 ppm, semi-tryptic peptides with up to one missed cleavage, static modifications were cysteine carboxyamidomethylation (+57.02 Da), lysine and N-terminal dimethylation (light formaldehyde 28.0313 Da; medium formaldehyde 32.0564 Da; heavy formaldehyde 36.0756 Da) and methionine oxidation and deamidation of asparagine and glutamine as variable modifications. A false discovery rate of 0.01 was applied both for spectrum-to-sequence matching and protein identification. A custom PERL script was used to filter semi-specific peptides and through database-lookup determine the corresponding prime and nonprime sequence (Demir et al 2022). Sequence logos showing substrate specificity were calculated as iceLogos using a local installation (Colaert et al. 2009).

### *In-planta* protein production and purification

C-terminally tagged Pip1 (pJK083; pL2M-P19-2x35S::Pip1-6xHis), Pip1(C153A/C154A) (pJK083; pL2M-P19-2x35S::Pip1(C153A/C154A)-6xHis), Rcr3 (pJK026; pL2M-P19-2x35S::Rcr3-6xHis), Rcr3(C153A/C154A) (pJK145; pL2M-P19-2x35S::Rcr3(C153A/C154A)-6xHis), Pip1+ (pJK674; pL2M-P19-2x35S::Pip1+-6xHis), Pip1+(C153A/C154A) (pJK675; pL2M-P19-2x35S::Pip1+(C153A/C154A)-6xHis) and Pip1++ (pJK676; pL2M-P19-2x35S::Pip1++-6xHis) were produced and purified from *N. benthamiana* as previously described (Schuster et al., 2022). Briefly, proteins were transiently produced by agroinfiltration as described above. Six days-post-infiltration AF was extracted as described above. Protein purification was performed at 4°C to prevent protease self-degradation. 250 μL Ni-NTA resin (Qiagen) was equilibrated with 20 column volumes (CV) ice-cold purification buffer (50 mM Tris-HCl, pH 7.4, 150 mM NaCl) in a small gravity column (Mini bio spin, Biorad). 8 to 10 mL apoplastic fluid containing the His-tagged proteins was loaded twice onto the resin. The Ni-NTA agarose was washed with 10 column volumes washing buffer (50 mM Tris-HCl, 150 mM NaCl, 10 mM imidazole, pH 8) and the protein was eluted using elution buffer (50 mM Tris-HCl, 150 mM NaCl, 50 mM imidazole, pH 8). Finally, the purified protein was concentrated by centrifugation in 3 kDa cut-off filters (Amicon Ultra-15 Centrifugal Filter Units, Merck) at 4,500 x g, 4°C, flash-frozen in liquid nitrogen and stored at -80°C. Protein purity was verified by running the protein on 15% SDS-PAGE gels followed by Coomassie staining.

### Substrate specificity assay

1µg pure protein was incubated in reaction buffer (50 mM sodium acetate, pH 5.0, 10mM DTT) supplemented with either 4 mM FR-AMC (Sigma) or LR-AMC (Bachem) substrate. Fluorescence was measured using a 96 well plate reader (Tecan Infinite M200) every 20 seconds for 2 hours at ex360/em460 and the 1-hour timepoint was selected for data visualization. Data points were analysed based on detected total changes in emission, relative to t0.

## ACKNOWLEDGEMENTS

We thank Urszula Pyzio for excellent plant care; Sarah Rodgers, Caroline O’Brian and Patricia Bowman for technical assistance; John Baker for photography; Sylvestre Marillonnet and Nicola Patron for providing plasmids via AddGene.

## AUTHOR CONTRIBUTIONS

JK generated most of the dataset by cloning, protease expression, ABPP and HR analysis; MS purified immune proteases and performed assays with fluorescent dipeptides; FD and PH performed PICS analysis on purified proteases provided by JK; OM, SK, PK, RO, SR, AB and BM supported the analysis of the mutant proteases by transient expression and/or ABPP and/or HR assays; JK, SK and RH conceived the project; and JK and RH wrote the article with input from all co-authors.

## SUPPLEMENTARY INFORMATION

**Figure S1** *Nb*Rcr3(G184N+DPS) induces Avr2/Cf-2-dependent HR.

**Figure S2** 33 Pip1/Rcr3 hybrids are active proteases.

**Figure S3** 30 Pip1/Rcr3 hybrids can be inhibited by Avr2.

**Figure S4** Eight Pip1/Rcr3 hybrids trigger Avr2/Cf-2-dependent HR.

**Figure S5** Selection of Rcr3-specific residues that may contribute to HR.

**Table S1:** Part-3 tested hybrids and results.

**Table S2:** Part-5 tested hybrids and results.

**Table S3:** Part-6 tested hybrids and results.

**Table S4:** Plasmids used in this study.

**Supplemental dataset S1:** Uncropped gels.

**Supplemental dataset S2:** Plasmid maps.

**Supplemental dataset S3:** Scripts, raw data, and statistics for HR quantification.

