## Supplementary figures and images for "Bioengineering secreted proteases converts divergent Rcr3 orthologs and paralogs into extracellular immune co-receptors"

### BestHR_Hybrids.pdf

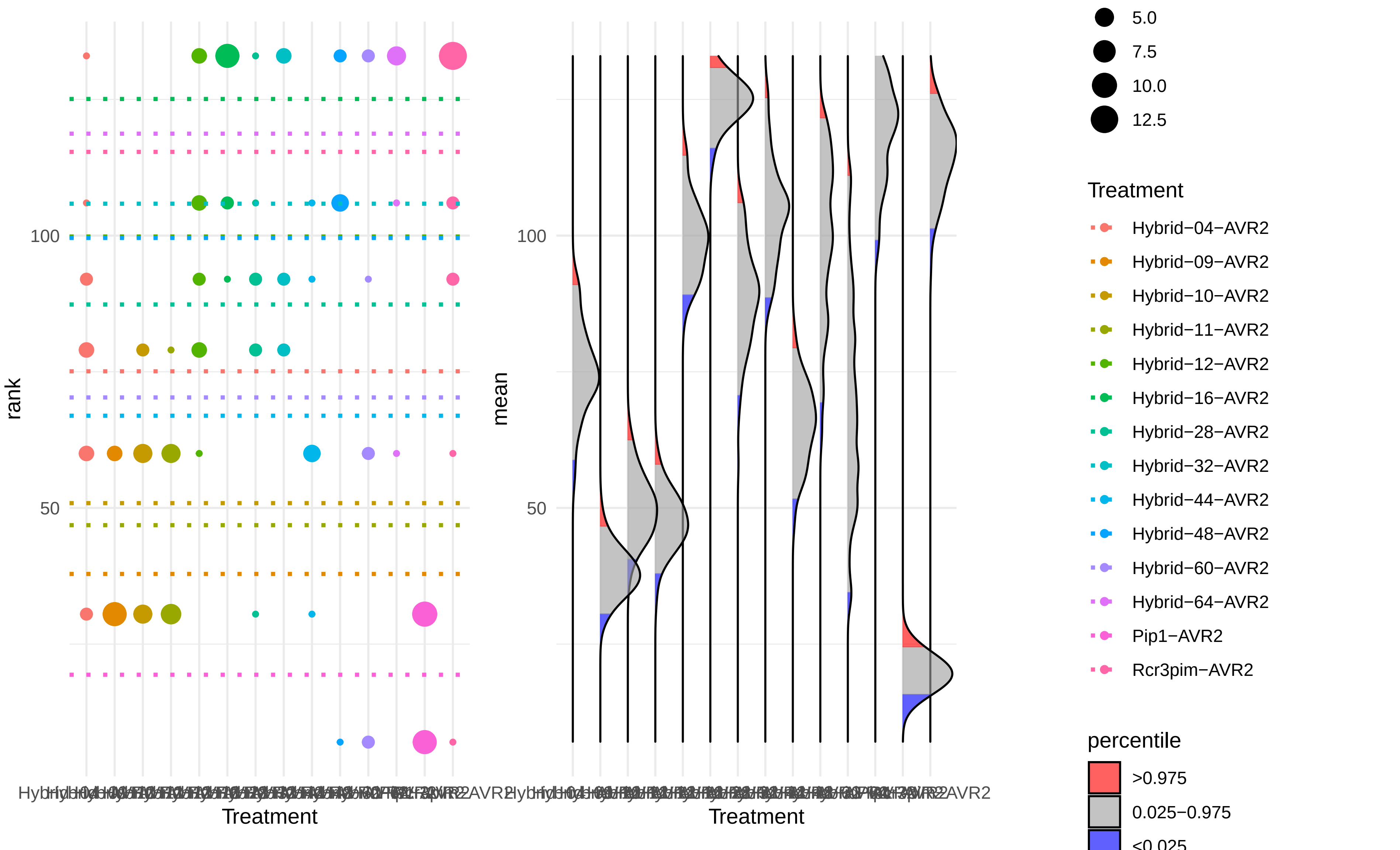

### BestHR_NbRCR3.pdf

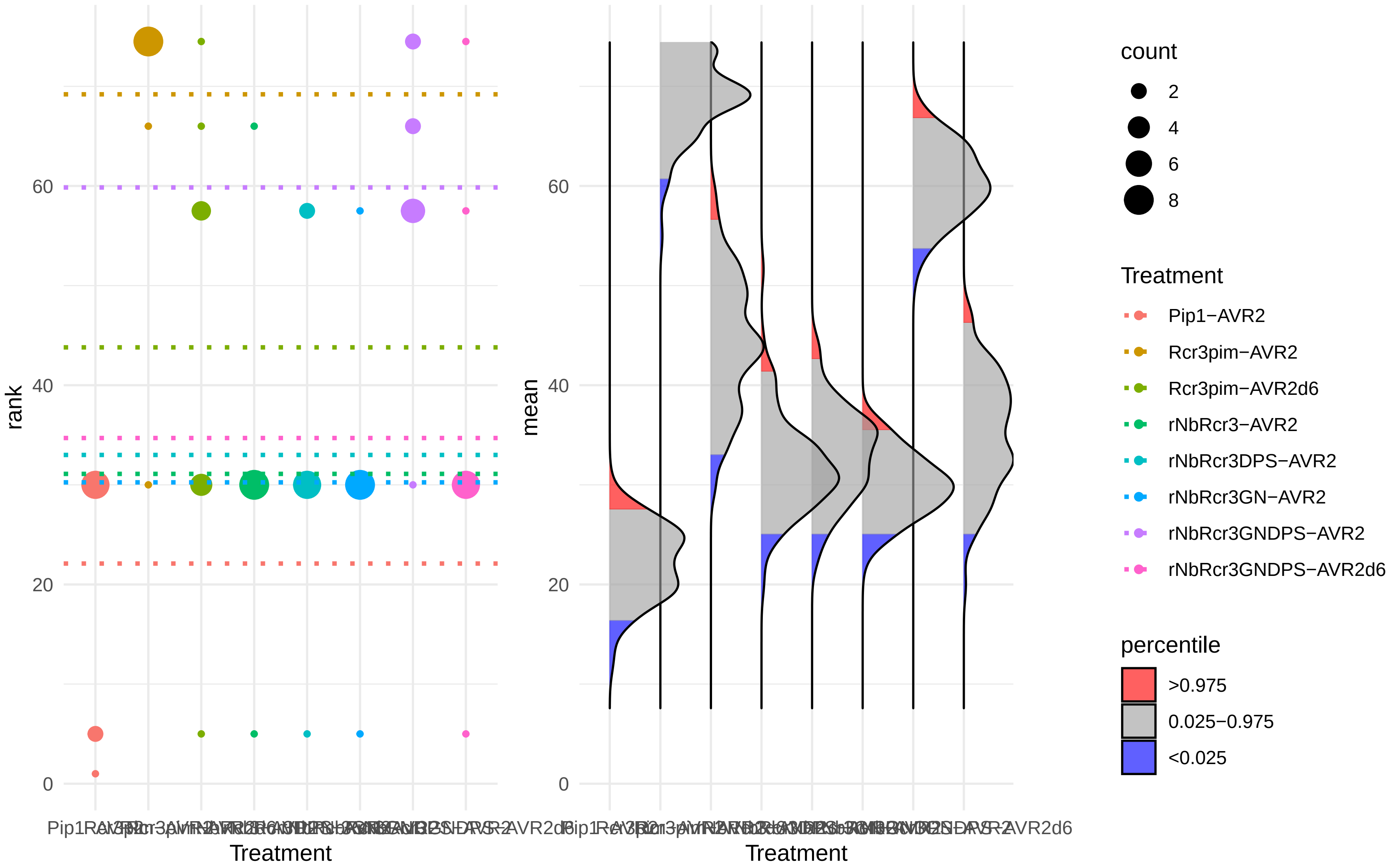

### BestHR_NtRCR3.pdf

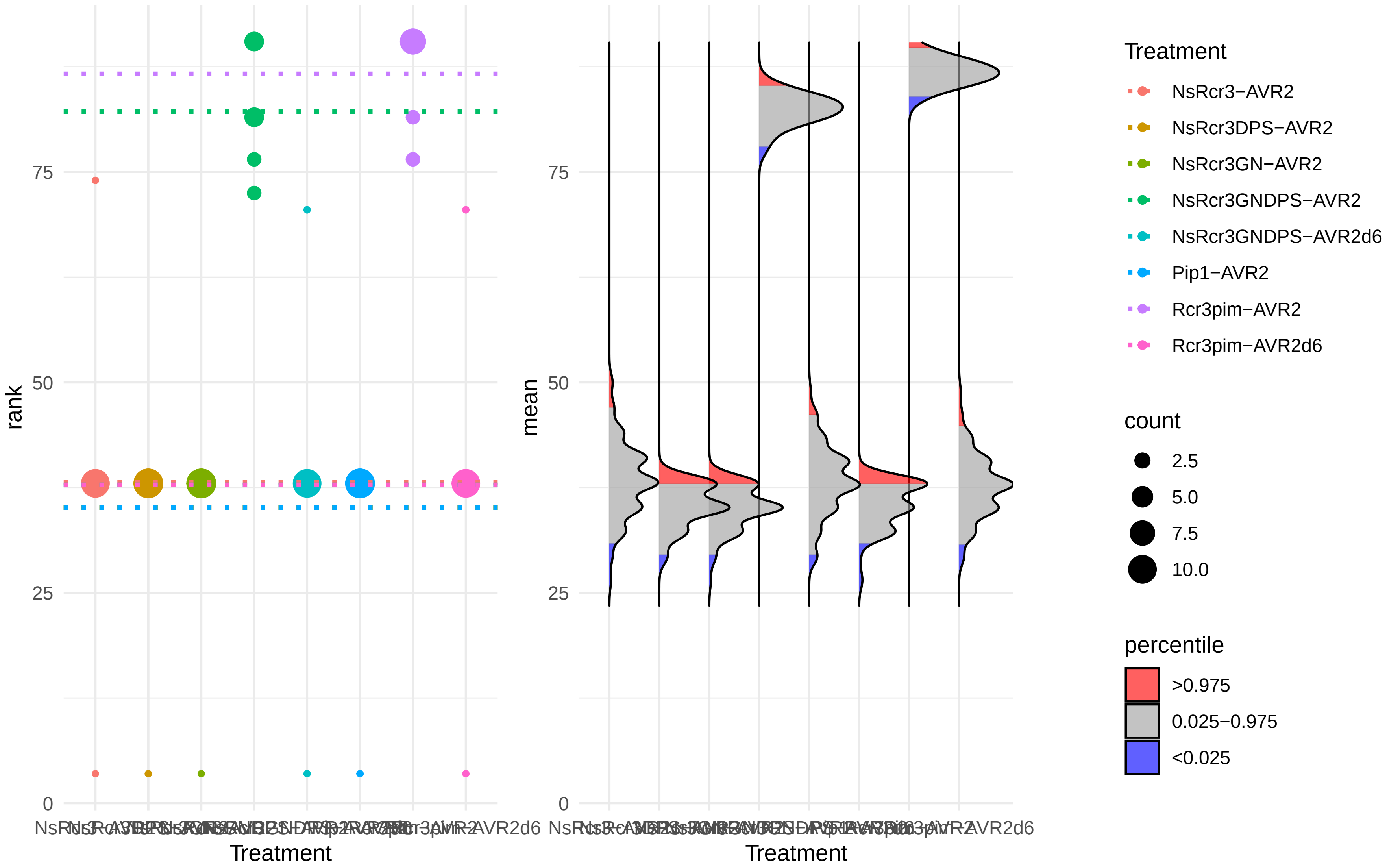

### BestHR_Pip1plus.pdf

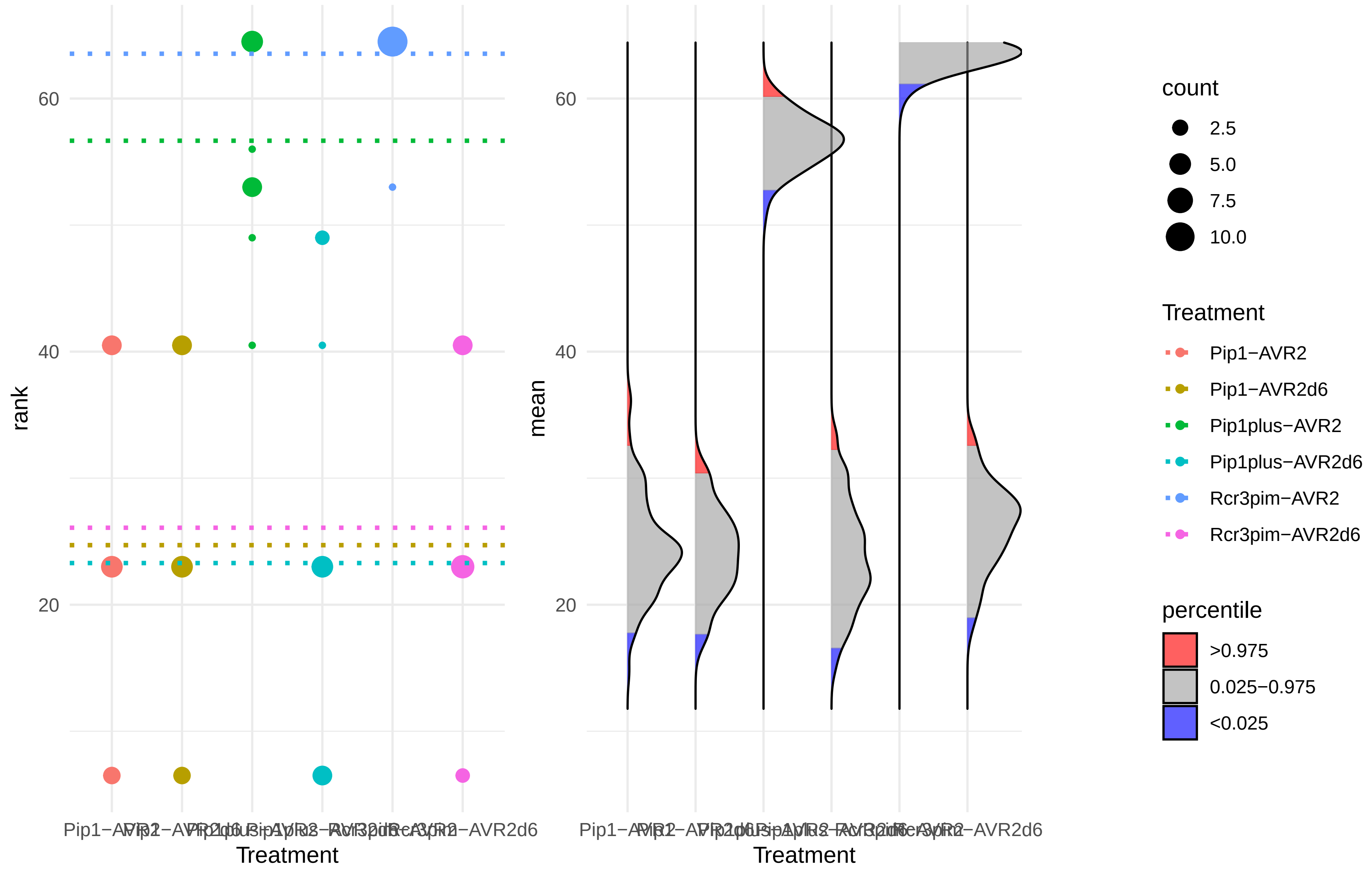

### BestHR_Pip1plusplus.pdf

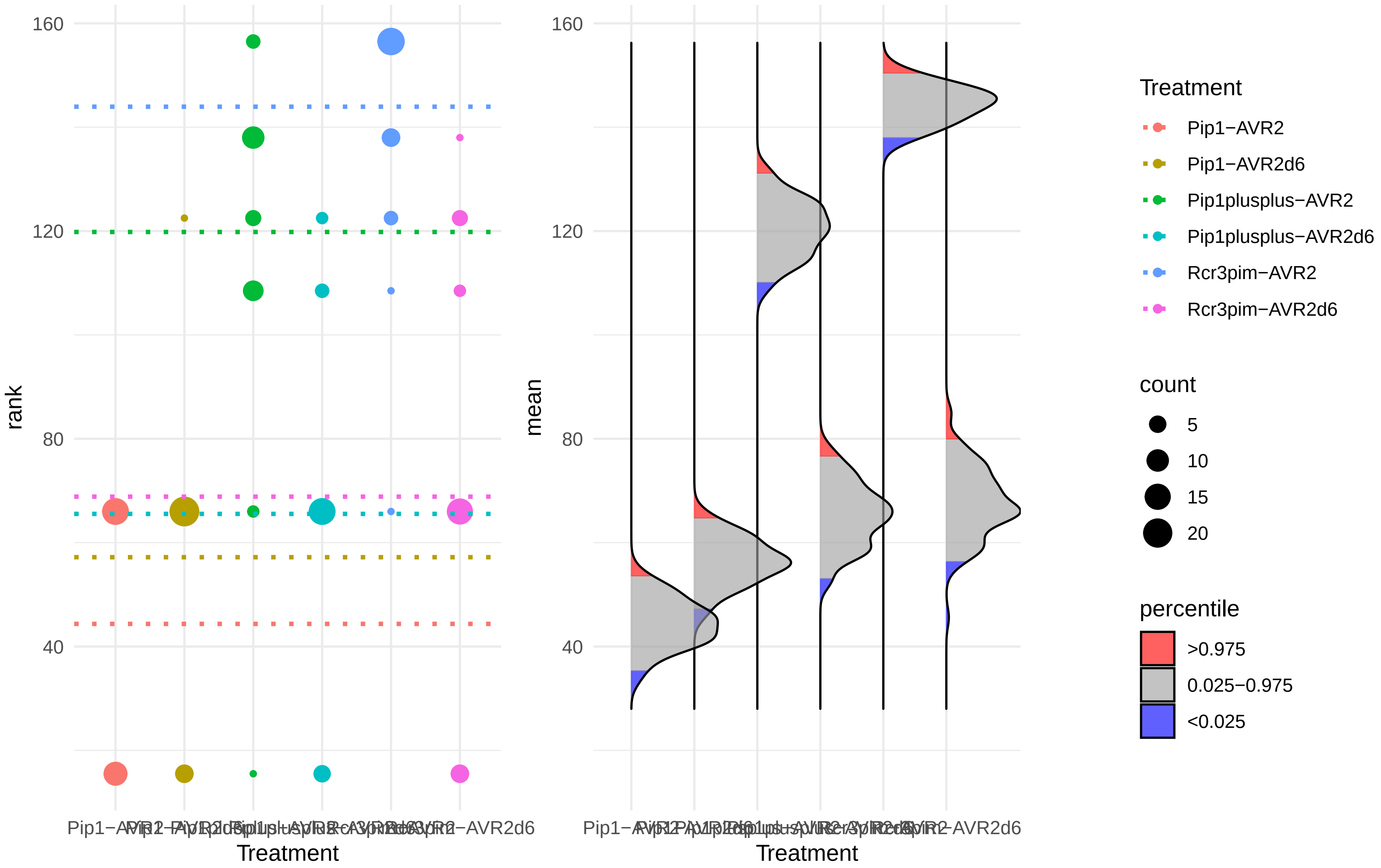

### BestHR_pJK235_03.pdf

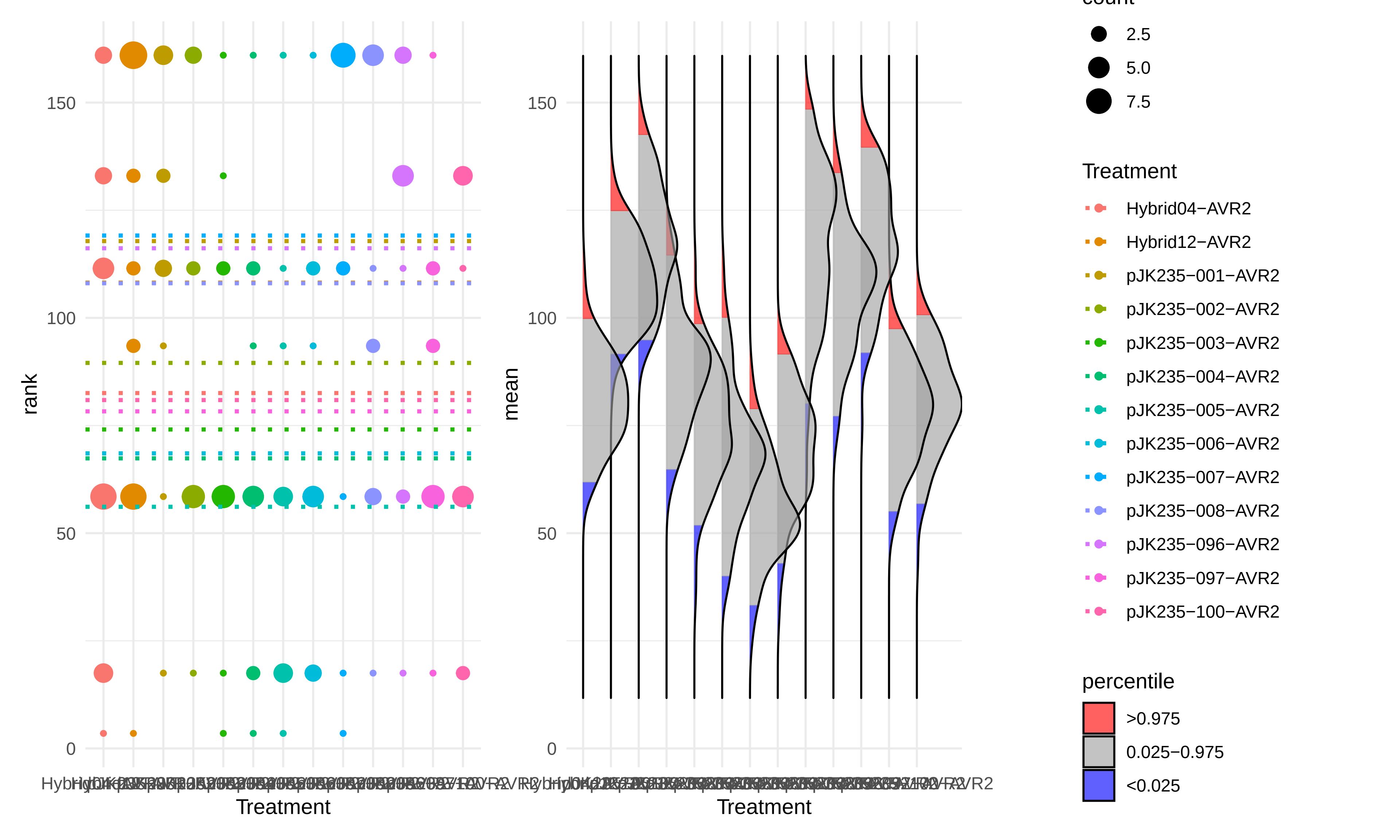

### BestHR_pJK235_05.pdf

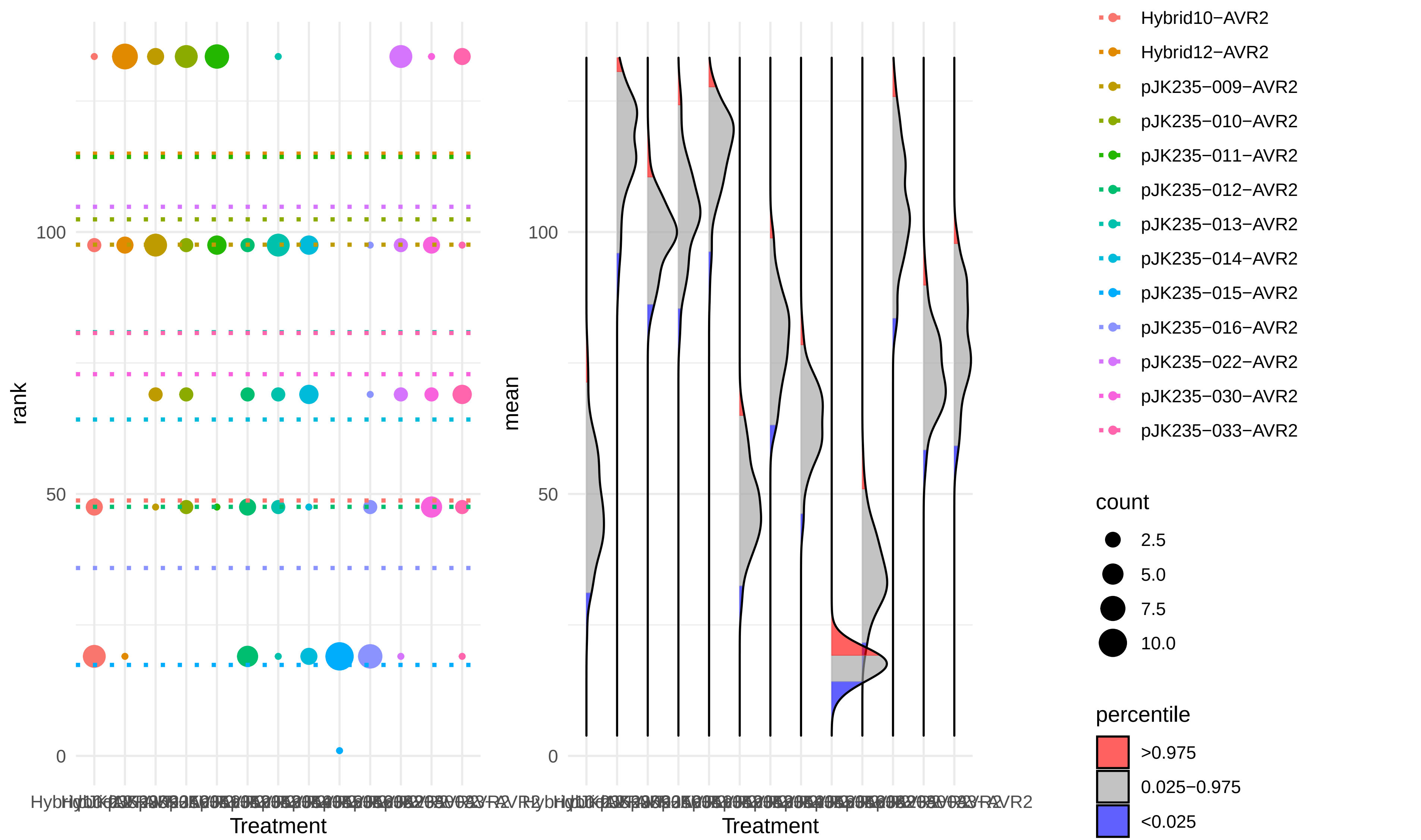

### BestHR_pJK235_06.pdf

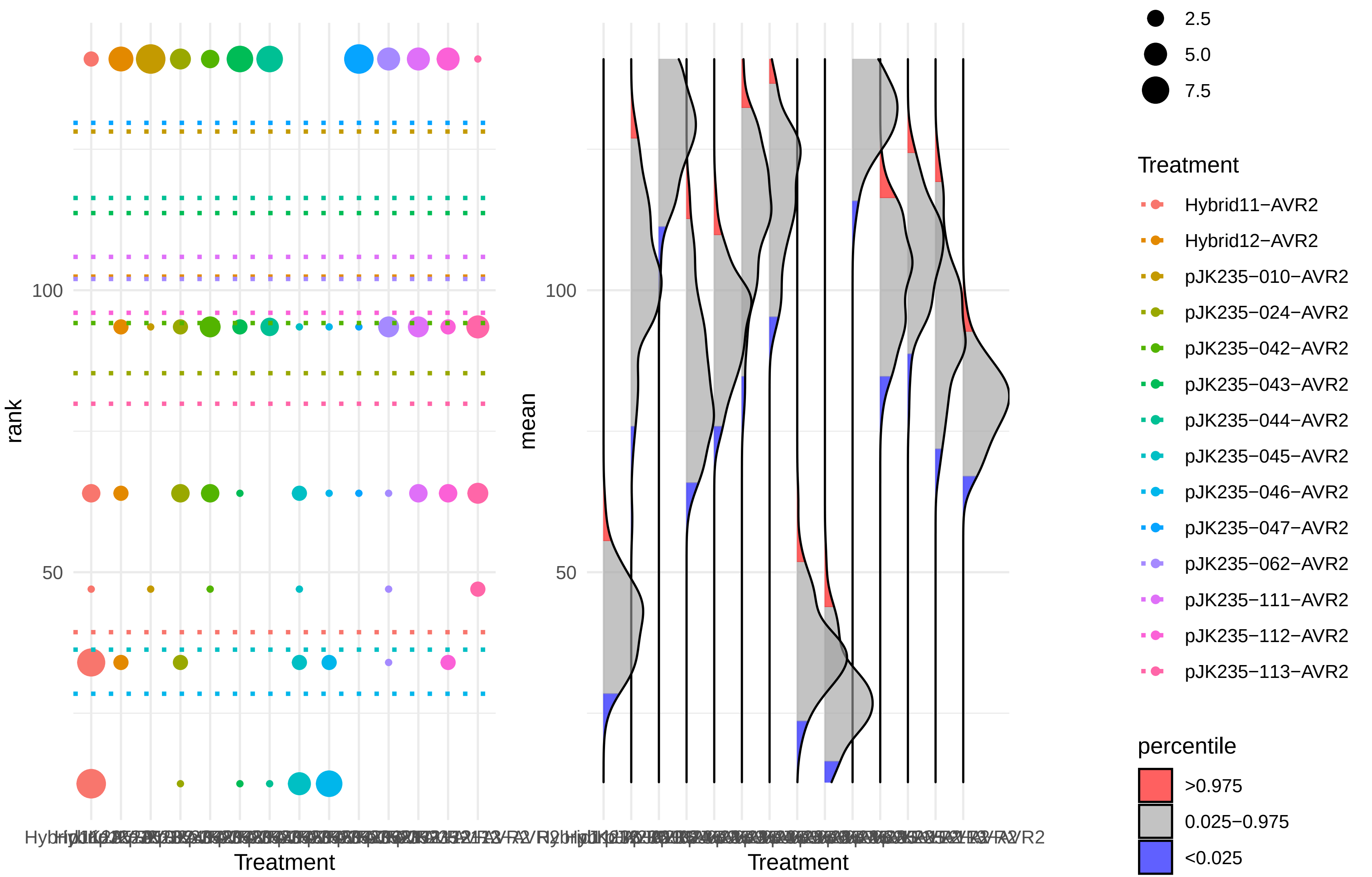

### BestHR_SmRCR3.pdf

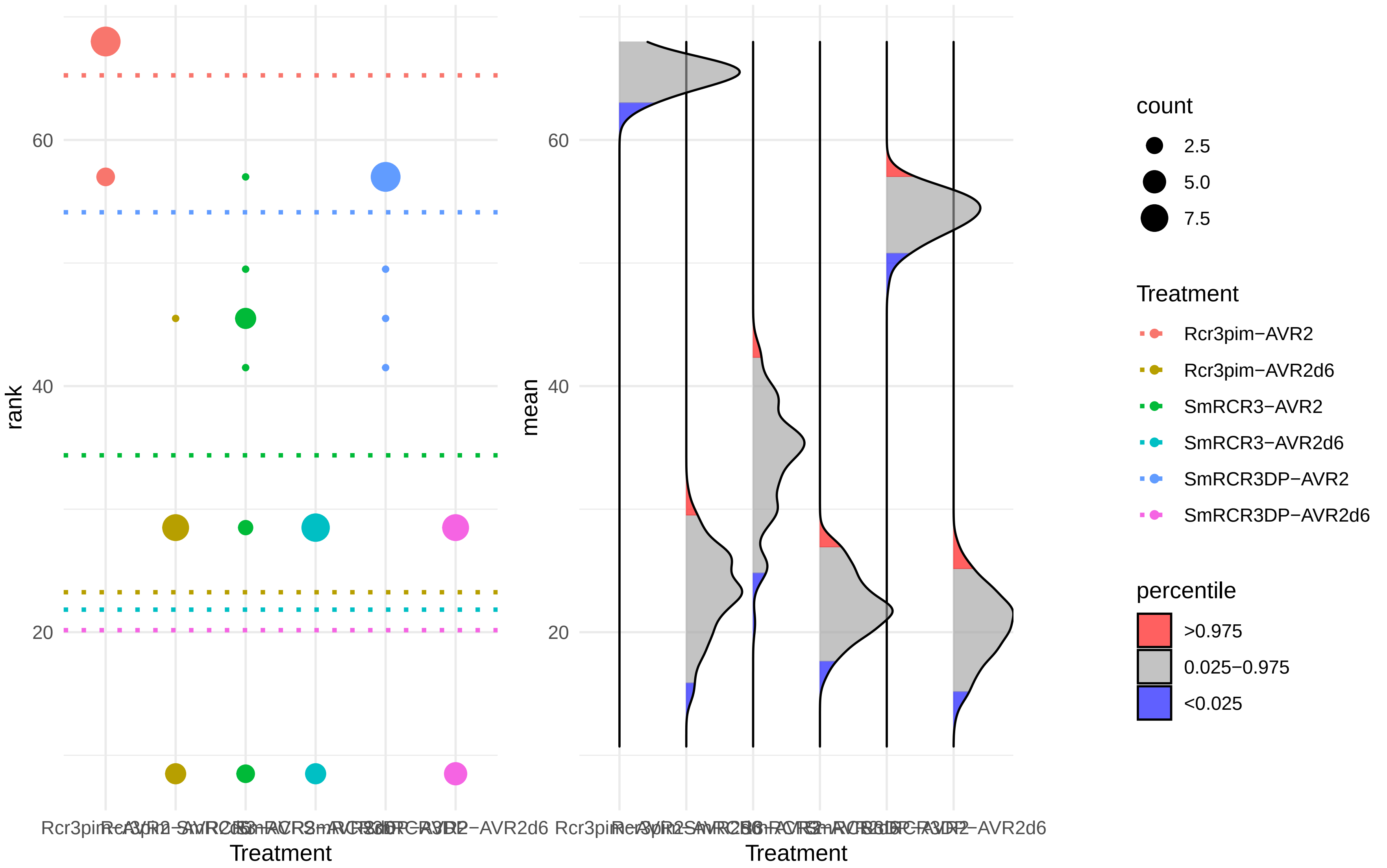

### BestHR_SmRCR3_fragment.pdf

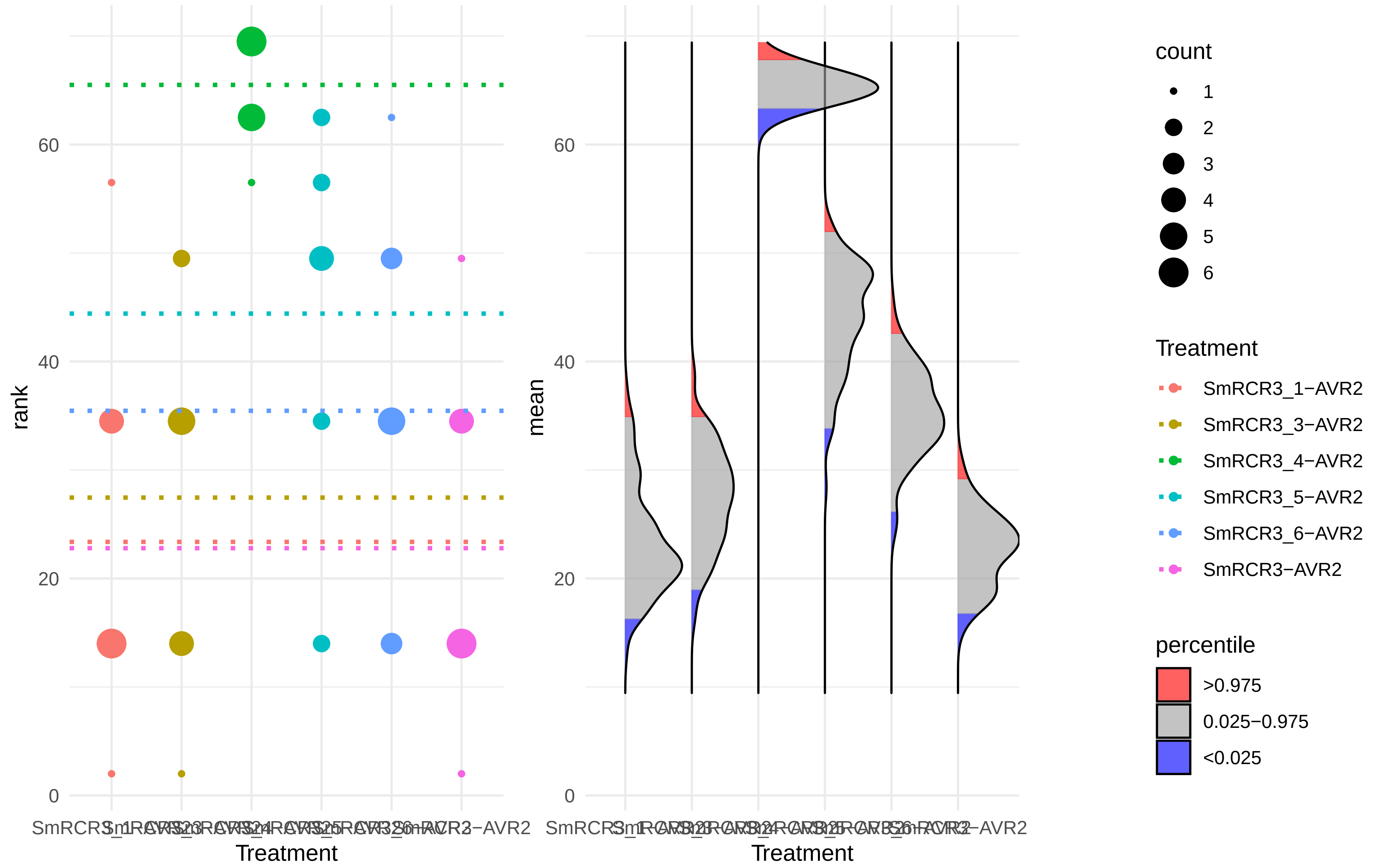

### Dataset S1

Figure 1E)

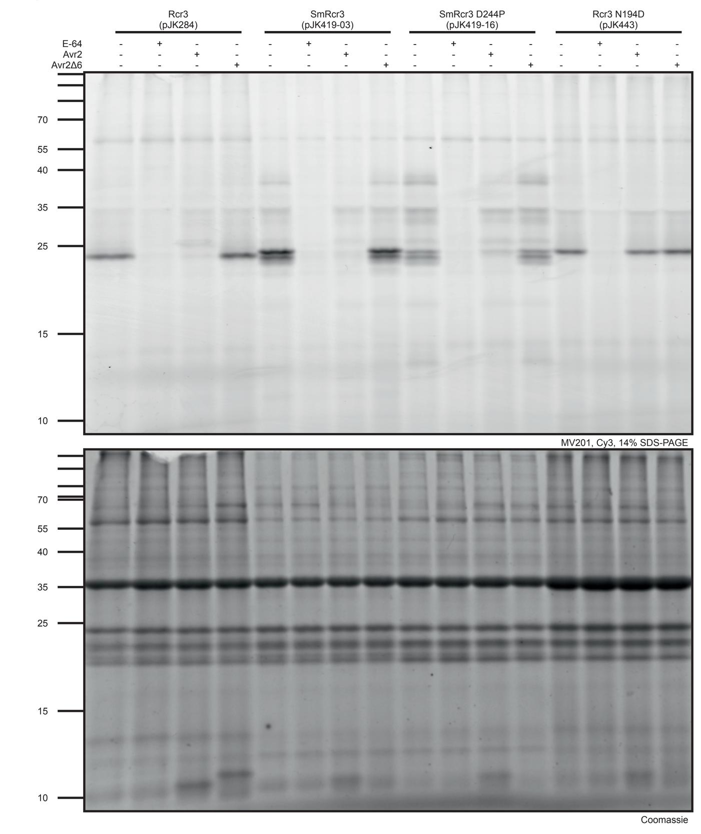


Figure 2B)

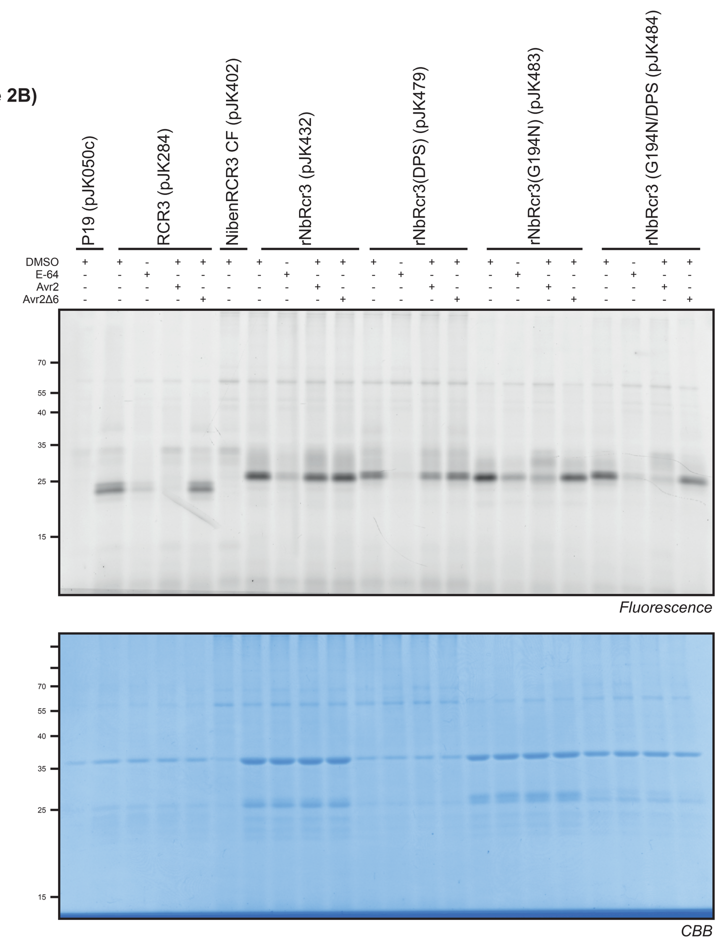


Figure 2C)

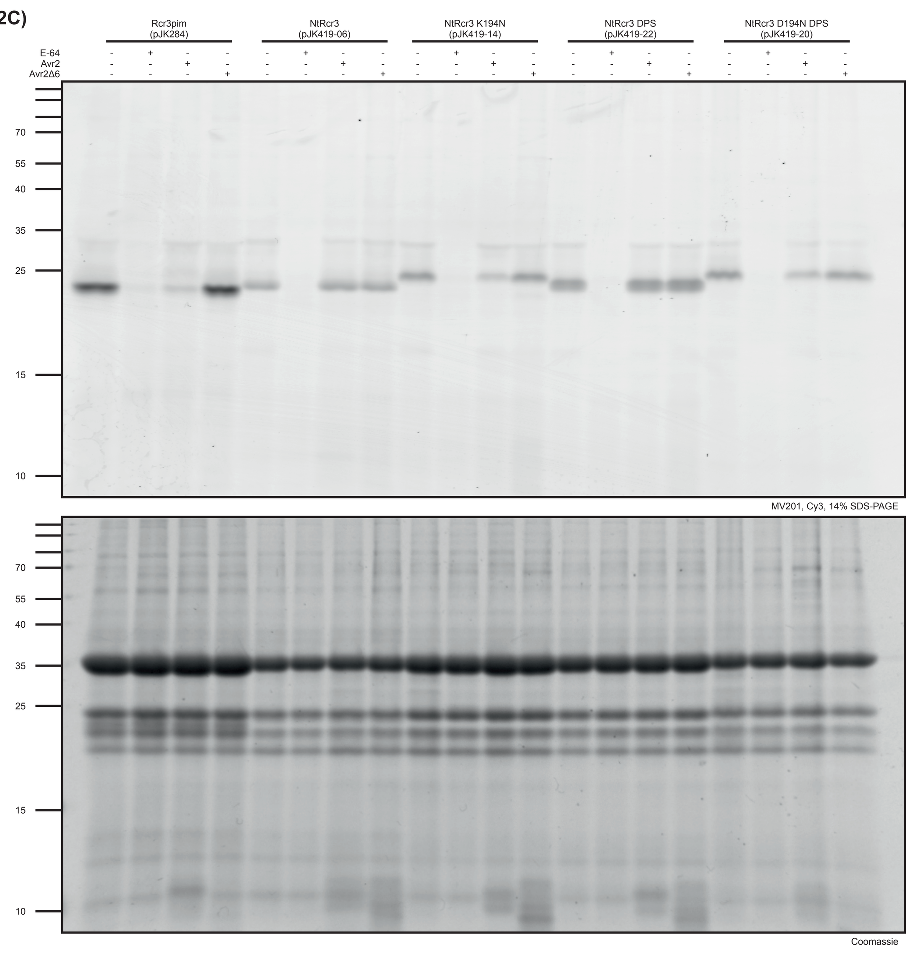


Figure 3C)

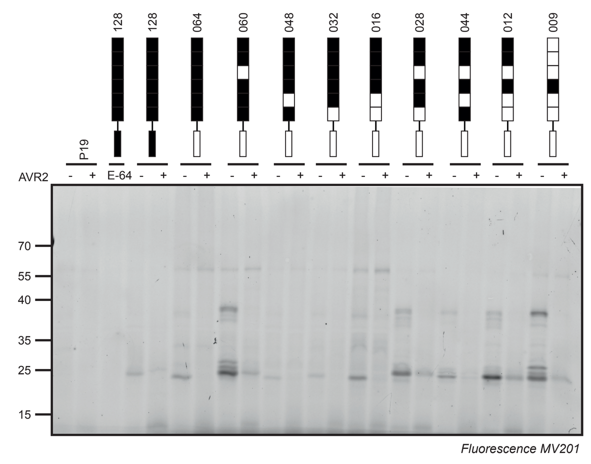


Figure S1)

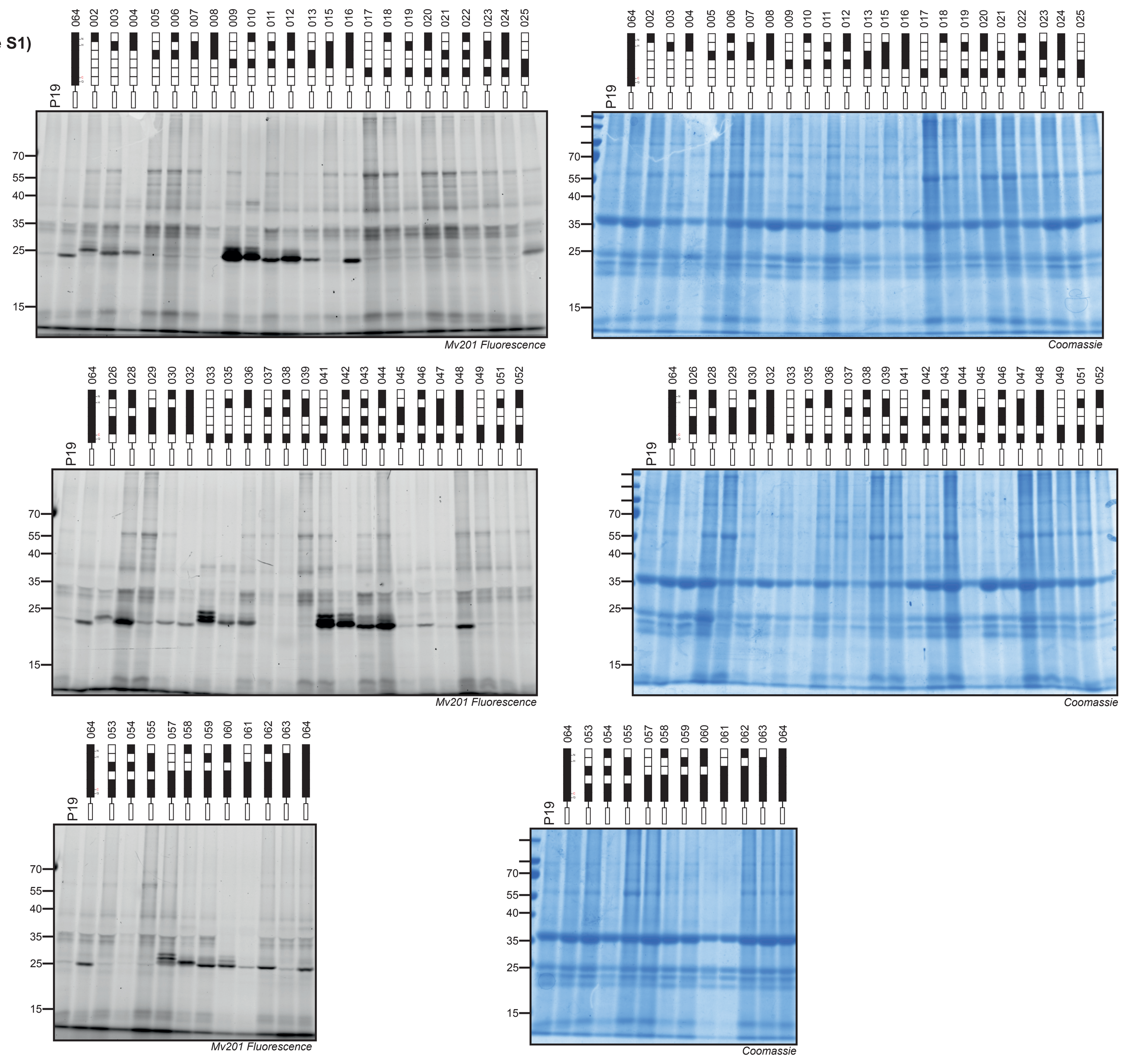


Figure S2)

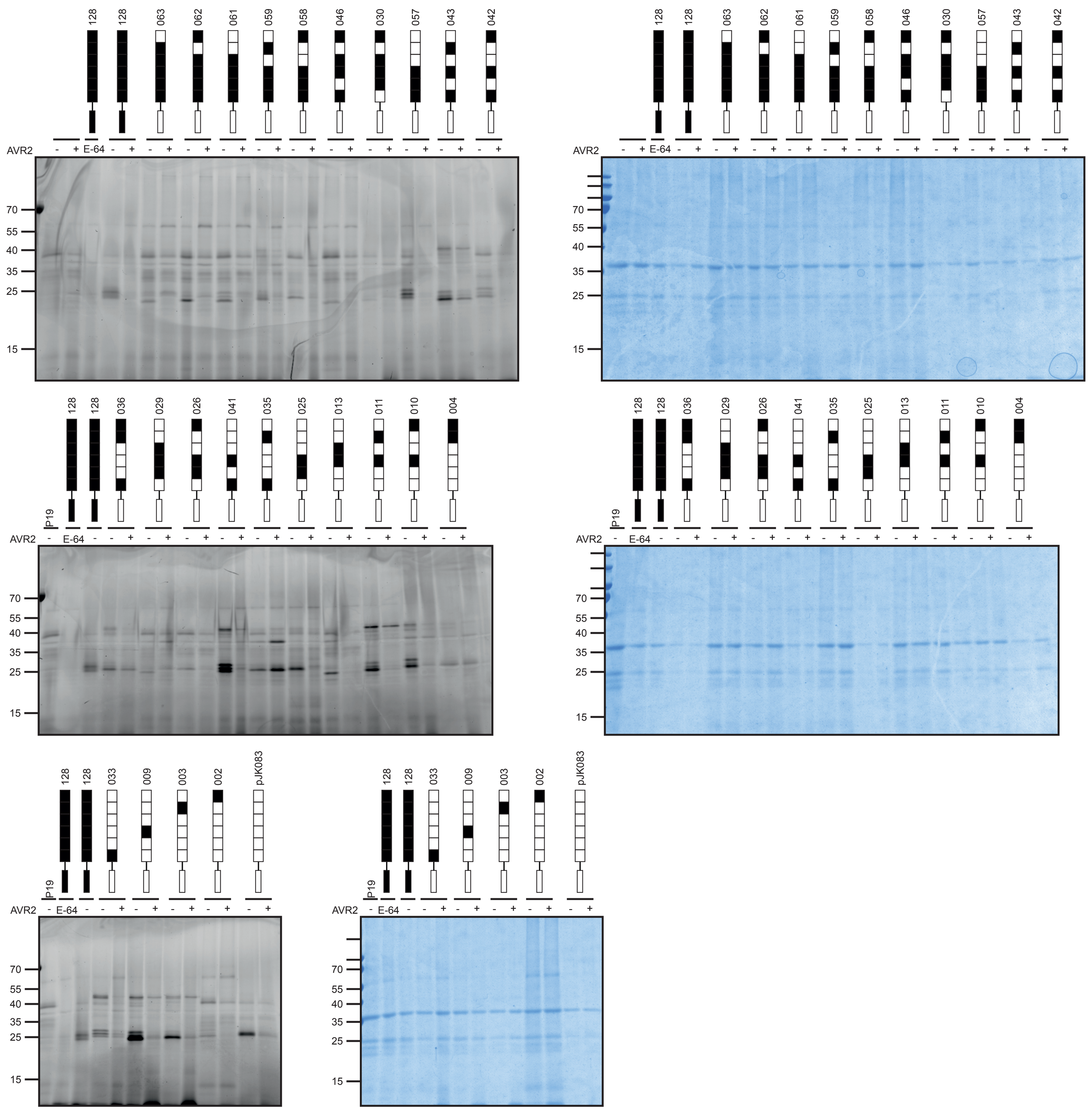
