## Supplementary material for "Bioengineering secreted proteases converts divergent Rcr3 orthologs and paralogs into extracellular immune co-receptors": Dataset S3: Hybrids_HR_quantification.R

```
# Packages

library(tidyverse)
library(besthr)

################################################################
################################################################
################################################################

# Read in data

df <- readxl::read_excel("Hybrids_HR_quantification.xlsx") %>%
  filter(!is.na(Score)) %>% # Remove missing values
  mutate(Replicate = factor(Replicate, levels = c("1")),
         Genotype = factor(Genotype, levels = c("wt")),
         Receptor = factor(Receptor, levels = c("Cf2")),
         Protease = factor(Protease, levels = c("Rcr3pim", "Pip1", "Hybrid-64", "Hybrid-60", "Hybrid-48", "Hybrid-44", "Hybrid-32", "Hybrid-28", "Hybrid-16", "Hybrid-12", "Hybrid-11", "Hybrid-10", "Hybrid-09", "Hybrid-04")),
         Effector = factor(Effector, levels = c("AVR2", "AVR2d6")),
         Score = factor(Score, levels = c("0", "1", "2", "3", "4", "5", "6", "7")),
         Treatment = paste0(Protease, "-", Effector))

# Group data by Score, NLR1, NLR2, Effector, and count number of occurrences 
df_grouped <- df %>% 
  group_by(Score, Receptor, Protease, Effector, Treatment) %>%
  summarise(count = n())

################################################################
################################################################
################################################################

# Assign colours
dfcols <- c("AVR2" = "#8faad0",
            "AVR2d6" = "#c9a9ff")

# Plot setup with Effector on x-axis and Score on y-axis
df_plot <- ggplot(df_grouped, aes(x = Treatment, y = Score)) +
  # Add a dotplot where the size and fill of dots are determined by count and Effector
  geom_count(
    aes(size = count, fill = Effector), 
    shape = 21, 
    stroke = 1,
    show.legend = TRUE
  ) +
  # Overlay original data as jittered points colored by Replicate
  geom_jitter(
    data = df, 
    aes(color = Replicate), 
    size = 0.5, 
    width = 0.3, 
    height = 0.2
  ) +
  # Define manual fill colors, set y-axis limits, and add labels
  scale_fill_manual("", values = dfcols, breaks = NULL) +
  scale_y_discrete(limits = c("0", "1", "2", "3", "4", "5", "6", "7")) +
  labs(x = "", y = "Cell death index", size = "Count") +
  # Define manual colors for different replicates and the size of the dots
  scale_color_manual(values = c ("#d0800b", "#00aa00","#F8F300")) +
  scale_size_continuous(range = c(1, 12)) +
  #facet_wrap( ~ Protease, nrow = 1, scales = "free_x") +
  theme_bw() +
  theme(
    axis.text = element_text(size = 10, color = "black"),
    axis.title = element_text(size = 10, color = "black"),
    axis.text.x = element_text(angle = 90, hjust = 1, vjust = 0.5),
    legend.title = element_text(size = 10),
    legend.text = element_text(size = 10),
    strip.text.x  = element_text(size = 10, colour = "black"),
    strip.background = element_blank(), 
    strip.placement = "outside",
    legend.key.size = unit(0.35,'cm')
  )


# Display the plot
df_plot

ggsave("Quantification_Hybrids.pdf", width = 10, height = 6)

################################################################
################################################################
################################################################

hr_est_combined_Sample <- estimate(df, Score, Treatment, Replicate, Control = "Rcr3pim-AVR2")

plot(hr_est_combined_Sample)

ggsave("BestHR_Hybrids.pdf", width = 10, height = 6)
```
