## Supplementary material for "Bioengineering secreted proteases converts divergent Rcr3 orthologs and paralogs into extracellular immune co-receptors": Figure S1

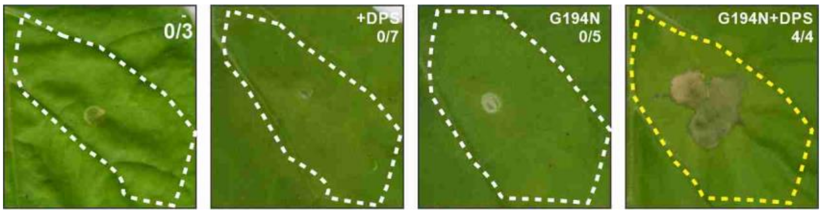


**Figure S1** r*Nb*Rcr3(G184N+DPS) induces Avr2/Cf-2-dependent HR.

Resurrected (r) *Nb*Rcr3 and derived +DPS and G194N mutants were co-expressed with Avr2 and Cf-2 by agroinfiltration of *Nicotiana benthamiana* with OD = 0.25 each in a 1:1:1 ratio. Images were taken at 5dpi. Numbers indicate the number of agroinfiltrated sectors showing HR symptoms.
